# An Empirical Bayes approach for the study of phenotypic evolution from high-dimensional data

**DOI:** 10.64898/2026.03.13.711658

**Authors:** Paola Montoya, Anne-Claire Fabre, Anjali Goswami, Helene Morlon, Julien Clavel

## Abstract

Multivariate phylogenetic comparative methods for modelling high-dimensional traits such as 3D shapes or gene expression profiles have been recently developed. However, these approaches are impractical and almost impossible to use when the number of traits exceeds a few thousands, as they become computationally prohibitive. We overcome these limitations by proposing a new maximum likelihood approach based on the Empirical Bayes framework. This approach takes into account the information of the complete covariances (among species and traits) to infer parameters and compare models of trait evolution for high-dimensional datasets. Through simulations, we demonstrate that the proposed approach can accurately estimate parameters of various trait evolution models, even when the number of traits is ten times larger than the number of lineages; it requires less memory and is at least 10 times faster than currently available approaches. This fast, efficient framework enabled us to extend the high-dimensional multivariate phylogenetic comparative toolkit by including an Ornstein-Uhlenbeck process with multiple optima to study adaptation to various selective regimes. Applying our approach to the evolution of jaw morphology in relation to dietary adaptation in mammals, we demonstrate morphological convergence in carnivorous and herbivorous lineages. The proposed Empirical Bayes framework, implemented in the R package mvMORPH, enables phylogenetic comparative methods to efficiently handle high-dimensional datasets and complex models of trait evolution.

---

Recent technological advances have enabled a focused effort on high-throughput phenotyping of quantitative traits, with the premise of better capturing and characterising phenotypic complexity and diversity (Davies et al. 2017; Goswami et al. 2019; Krassowski et al. 2020; Snead and Clark 2022; Ford et al. 2023; Blackburn et al. 2024; Goswami and Clavel 2025). These efforts are materialised in a growing number of databases containing thousands of high-resolution datasets. For instance, multiple anatomical structures are described through 3D images of museum specimens from a wide range of species, both extant and extinct (e.g., Phenome10k (Goswami 2015); MorphoSource (morphosource.org); MorphoMuseuM (Lebrun and Orliac 2016), DigiMorph (digimorph.org)), and usually analysed through geometric morphometric techniques. Similarly, molecular and genome-scale continuous traits such as gene expression profiles for multiple tissues across different taxa are becoming increasingly common (e.g., GEO (ncbi.nlm.nih.gov/geo/); Bgee (Bastian et al. 2025)). These comparative datasets hold significant potential to address a broad array of questions in biology because of their scale and detailed resolution. However, they remain challenging to analyse within a phylogenetic framework, not only because of their large sizes, but more importantly because the number of traits *p* often exceeds the number of species *n*, which prevents the use of most multivariate phylogenetic comparative methods (PCMs) based on likelihoods (Goolsby 2016; Adams and Collyer 2018; Clavel et al. 2019). Indeed, in these high-dimensional conditions (*p* >= *n*), the matrix describing the variance-covariances of the traits becomes singular. In practical terms, this means that this matrix is not invertible and we cannot estimate its determinant, two mathematical operations which are required to calculate the likelihood of all the classical models of multivariate trait evolution (Revell and Collar 2009; Bartoszek et al. 2012; Clavel et al. 2015; Goolsby et al. 2017). Various strategies have been employed to comply with the low-dimensionality requirements of standard PCMs. These include assuming independence between variables and ignoring covariance between traits, or using dimension reduction techniques such as principal component analysis (PCA) to obtain a reduced set of variables. However, these practices often lead to inaccuracies in both model selection and parameter estimation (Uyeda et al. 2015; Clavel and Morlon 2020).

Over the last decade, different PCMs directly applicable to high-dimensional data sets have been developed. For example, some methods estimate evolutionary rates and perform statistical analyses based on all pairwise trait (Euclidean) distances between species, rather than on the covariance matrix, thereby avoiding computation of the multivariate likelihood (Adams 2014a, 2014b, 2014c; Adams and Collyer 2015). However, these distance-based approaches are restricted to the Brownian motion model and ignore the trait correlations, which are precisely one of the key aspects that multivariate approaches aim to capture (Goolsby 2016; Clavel et al. 2019; Clavel and Morlon 2020). Goolsby (2016) proposed a pairwise composite likelihood (PCL) approach, where the problem is reduced to the evaluation of the likelihoods for all the possible pairwise combinations of traits (i.e., bivariate models ensuring that *p<<n*). Although the PCL method is both efficient and accurate (Goolsby 2016; Clavel et al. 2019), it does not directly provide an estimator of the full covariance matrix across all traits (Adams and Collyer 2018). Additionally, the inference made by the PCL method is not rotation-invariant (Adams and Collyer 2018), which makes it unsuitable for analysing datasets with no natural orientation, such as those obtained using landmark-based geometric morphometric techniques (Rohlf 1999). This type of data requires rotation-invariant approaches, in which the parameter inference is independent of the arbitrary orientation of the shapes under study. Finally, penalized likelihood (PL) techniques have been used to tackle the problem by obtaining improved and invertible estimators of the trait covariance matrix under different multivariate trait evolution models, through regularisation (Clavel et al. 2019). Regularisation (i.e., reducing the variability or constraining the parameters estimation) is an effective way to treat high-dimensional covariance matrices, commonly applied in the statistical literature (e.g., Engl et al. 2000; Daniels and Kass 2001; Bickel and Levina 2008; Calvetti and Somersalo 2018). In the PL approach the regularisation is performed by amending a penalty term to the multivariate likelihood with the objective to enforce the shrinkage of the covariance matrix estimators towards predefined, simpler, and well-behaved variance-covariance structures, overcoming most of the above-mentioned limitations. The approach has already successfully been applied to various high-dimensional datasets (e.g., Eliason et al. 2020; Fabre et al. 2021; Coombs et al. 2022; Goswami et al. 2022; Sansalone et al. 2024). However, PL-based approaches are computationally demanding as they require cross-validation (CV) techniques, which involve the repeated estimation and inversion of potentially large covariance matrices to infer the parameter controlling the level of penalisation (up to *n* times for the leave-one-out CV (LOOCV), see details in Clavel et al. (2019)). Although some algorithms were proposed to reduce the computational cost of the cross-validation procedure (Clavel et al. 2019), these computational tricks still scale heavily with *n* and *p*, and are limited to specific models (for instance they cannot be used on the high-dimensional multiple optima Ornstein-Uhlenbeck model we describe below). In practice, this computational cost limits the application of PL-based PCMs when datasets are very large (e.g., with thousands of traits).

In a Bayesian framework, covariance matrix regularisation is usually achieved by selecting an appropriate prior to concentrate the posterior probability distribution to a space of well-behaved covariance matrices. Although it was not directly used to handle high-dimensional cases, the Bayesian framework and specific priors were used to ensure, for instance, positive-definiteness of covariance matrices in previous phylogenetic studies (Lartillot and Poujol 2011; Cuevas et al. 2014; Caetano and Harmon 2017). In practice, the parameters of the model, including the covariance matrix, are directly sampled from the posterior distribution using computationally intensive Markov Chain Monte Carlo (MCMC) techniques (e.g., Lartillot and Poujol 2011; Cuevas et al. 2014; Höhna et al. 2016; Caetano and Harmon 2017), which would make them as prohibitive as PL on large datasets. Instead of this traditional Bayesian approach, an approximation known as Empirical Bayes or maximum marginal likelihood estimation can be used (Carlin and Louis 2000; Gelman et al. 2014). In the Empirical Bayes framework, prior parameters are estimated from the data, usually by maximum likelihood (Carlin and Louis 2000; Petrone et al. 2014). If appropriate conjugate priors are used, posterior distributions can then be computed rapidly using analytical solutions, rather than through intensive MCMC sampling, which can save substantial computational resources. The Empirical Bayes framework has been successfully applied to the regularisation of the covariance matrix problem in the multivariate normal case in contexts other than trait evolution (e.g., Haff 1980; Coluccia 2015). Usually, a Wishart or inverse Wishart conjugate prior is used, one parameter of which acts as an analogue to the parameter controlling the regularisation intensity in the PL approach (Efron and Morris 1976; Haff 1980; Sharma and Krishnamoorthy 1985; Champion 2003; Coluccia 2015).

Here, we propose a new and efficient phylogenetic comparative method to study phenotypic evolution from high-dimensional datasets in a strict multivariate framework (i.e., taking into account all the trait covariances), using the Empirical Bayes framework. The beginning of the Material and Methods section provides a brief overview of the general concepts of the proposed approach. We implement this approach in the function *mvgls()* from the R package mvMORPH (Clavel et al. 2015), publicly available on CRAN (https://cran.r-project.org/package=mvMORPH) and GitHub (https://github.com/JClavel/mvMORPH). Through simulations, we show that the method achieves accuracy comparable to current approaches, while significantly reducing the computational cost (at least 10 times faster, and with lower memory requirements). These improvements allow us to further extend the high-dimensional multivariate PCMs to complex models such as the Ornstein-Uhlenbeck process with multiple optima (Hansen 1997; Butler and King 2004; Beaulieu et al. 2012). We illustrate our approach by exploring the convergence of lower jaw (mandibles) shape in carnivorous and herbivorous mammals. Although multiple studies have shown morphological convergences related to diet, mainly in teeth and lower jaws traits, for specific mammal clades (e.g., Weller 1968; Hoshi 1971; Biknevicius et al. 1996; Pérez-Barbería and Gordon 1999; Figueirido et al. 2010; Price et al. 2012; Kim et al. 2016; Law et al. 2022; Wu 2022), few have addressed this question at the scale of all mammals. Among these studies, some have used detailed high-dimensional trait data from 3D-scan images. However, they have either not directly evaluated hypotheses related to adaptive convergence (Morales-García et al. 2021), not accounted for all covariances by treating the traits individually (Grossnickle 2020), or used a subset of PC axes (Fabre et al. 2021). The latter study identified notable similarities in lower jaw structure between marsupials (Metatheria) and placentals (Eutheria) mammals, which were attributed to ecomorphological convergences, but also identified a significant impact of early development on jaw morphology. Specifically, the short gestation and lengthy period of postnatal care, during which marsupials young suckle continuously for many months, which is significantly longer than most placentals other than Primates (Weisbecker and Goswami 2010), constrains the morphological evolution of their jaws (see also Lillegraven 1975; Werdelin 1987; Conith et al. 2022). Here, we use the Empirical Bayes approach to re-analyse the dataset of Fabre et al. (2021). Specifically, we investigate how the lower jaw evolved in response to selective pressures associated with adulthood dietary regimes on the one hand, and constraints such as those imposed by extended suckling during early development on the other hand.

## Materials and Methods

### General overview of the Empirical Bayes approach

Here, we present the Empirical Bayes approach, an efficient method for fitting and comparing multivariate models of trait evolution, applicable to high-dimensional trait datasets. Its current implementation allows to Bit five trait evolution models: the Brownian Motion (BM), Early Burst (EB), Pagel’s lambda, and Ornstein-Uhlenbeck (OU) models, including the Ornstein-Uhlenbeck process with multiple optima (OUM).

In brief, the Empirical Bayes approach addresses issues arising from high-dimensional datasets (where *p > n*), by using a Bayesian formulation of the penalisation strategy employed in previous studies (e.g. penalized likelihood; Clavel et al. 2019). By eliciting an appropriate prior distribution for the matrix describing the covariances between the traits, the likelihood can be numerically evaluated. An excellent prior distribution for this purpose is the inverse Wishart distribution. This is due to its conjugacy with the multivariate Gaussian distribution, which underlies most phylogenetic comparative models of trait evolution. Conjugacy makes possible to integrate over all possible covariance matrices compatible with that prior analytically, without relying on computationally intensive Markov Chain Monte Carlo (MCMC) techniques (see *The Empirical Bayes formulation*). This has three practical advantages over previously proposed approaches. Firstly, it alleviates the need to compute explicitly the costly covariance matrix when *p* is large, because it is integrated out analytically. Secondly, unlike alternative approaches such as penalised likelihood, it does not rely on computationally intensive iterative procedures, such as cross-validation, to estimate the level of penalisation required for a robust model Bit. These two properties are key to considerably reducing the computational burden suffered by previous approaches, making the approach scalable to large datasets. Thirdly, it is possible to obtain an estimator of the matrix describing the covariances between the traits *a posteriori* (see *Regularised estimator of the variance-covariance matrix **R***). This is convenient for using comparative approaches such as multivariate phylogenetic regressions, and MANOVA tests.

### The Empirical Bayes formulation

In what follows, we use bold capital letters to refer to matrices, regular lowercase letters for scalars, and subscripts for the dimensionality of the given elements. ***X***^−1^, ***X***^*T*^, |***X***| indicate the inverse, the transpose, and the determinant of the matrix ***X*** respectively, while *vec*(***X***), the vectorised version of matrix ***X***. Finally, ⊗ is the Kronecker product between two matrices. A summary of all the parameters used throughout the text is provided in the supplementary material (Table S1).

In the classical multivariate models of trait evolution, such as Brownian motion (BM), Ornstein-Uhlenbeck (OU), or Early Burst (EB) processes, the traits (*vec*(***Y***)) follow a multivariate normal distribution (𝒩) with mean *vec*(***Θ***) and variance-covariance ***V***_*np* *x* *np*_ (Revell and Harmon 2008; Bartoszek et al. 2012; Clavel et al. 2015; Goolsby et al. 2017). When ***V*** can be decomposed as the Kronecker product of the matrices ***C***_*n* *x* *n*_describing the phylogenetic variance-covariance among species for a given evolutionary model, and ***R***_*p* *x* *p*_ describing the variance-covariance among traits for the same model (***R*** ⊗ ***C***; see Revell and Harmon 2008; Clavel et al. 2015, 2019), then ***Y*** is said to follow a matrix-variate normal distribution (ℳ𝒩) and can be more compactly represented as:

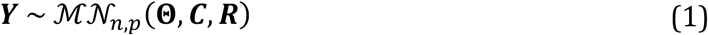

Note that equation (1) is equivalent to *vec*(***Y***) ∼ 𝒩(*vec*(***Θ***), ***V***), where ***V*** = ***R*** ⊗ ***C***. The mean ***Θ*** in equation (1) is given by the ancestral states (***β***) for each species and traits (equation 2). The parameters of the evolutionary model controlling the variance-covariance matrix ***C*** (*e.g.,* such as the selection strength ***α*** or the decay rate parameter *r* for Ornstein-Uhlenbeck or Early-Burst models respectively; Blomberg et al. 2003; Beaulieu et al. 2012; Clavel et al. 2015) are typically estimated by maximising the log-likelihood of the matrix-variate normal distribution in equation (1) with numerical methods, while closed-form solutions are used for the maximum likelihood estimates of ***Θ*** and ***R***. The maximum likelihood estimator for ***Θ*** is:

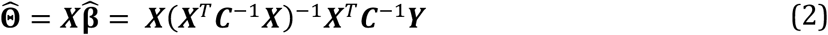

where ***X*** is a design matrix mapping the ancestral states ***β*** for each species and traits (a one-column matrix of ones, if a single ancestral state is estimated per traits). The maximum likelihood estimator for the variance-covariance matrix ***R*** is given by ***R̂*** = (***Y*** − ***Θ***)^*T*^***C***^−1^(***Y*** − ***Θ***)/*n* (Revell and Harmon 2008). However, in high-dimension, when *p > n*, this estimator becomes singular and prevents the evaluation of the log-likelihood as discussed earlier.

Following previous studies on Bayesian regularisation of the covariance matrix (e.g., Champion 2003; Coluccia 2015), we set an Inverse Wishart distribution (*W*^−1^(*k*, **ψ**)) prior on ***R***, a usual conjugate prior for the covariance matrix of a multivariate normal distribution (Gupta and Nagar 2000; Gelman et al. 2014), with degrees of freedom *κ* = *v* + *p* − 1 and a *p x p* positive definite scale matrix **ψ** = **Ψ**. Under this formalism, *v* is a parameter larger than 2 to fulfil the Inverse Wishart requirement of *κ* > *p* + 1 (Gupta and Nagar 2000). Using this conjugate prior to integrate ***R*** out of the equation (1), one obtains a marginal distribution for ***Y*** that follows a matrix-variate *T* distribution (Gupta and Nagar 2000; Iranmanesh et al. 2010):

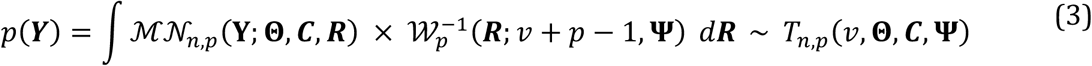

Here, we fixed *v* = *p* + 1, which results in a less diffuse prior than would be obtained by setting *v* to its minimum possible value. This ensures that the level of regularisation scales with the dimensionality of the dataset, a desirable property in regularisation as discussed in the next section. The choice of the scale matrix **Ψ** defines the shrinkage target (hereafter referred to as the “target”) of ***R*** and is typically a simpler well-behaved and invertible matrix. Like in the PL approaches, a natural candidate for this target matrix is a scaled identity matrix, such as **Ψ**_***iden***_ = *μ****I***_*p*_ (Coluccia 2015). An alternative target matrix can be defined as **Ψ**_***diag***_ = *μ****D***_*p*_, where ***D*** is a diagonal matrix with individual trait variance estimates (Clavel et al. 2019). The parameter *μ* scales uniformly the matrix **Ψ** and acts as an analogue to the parameter controlling the intensity of the regularisation in PL. These two targets trade-off between better robustness of the regularisation with respect to the structure in the data (**Ψ**_***diag***_; e.g., highly heteroscedastic datasets) and rotation invariance (**Ψ**_***iden***_), a requirement for datasets with no natural orientation like those from landmark-based geometric morphometrics (Rohlf 1999). Unlike the classical penalised likelihood-based approaches, which require a costly cross-validation procedure, the Empirical Bayes formulation allows us to infer the parameter *μ* directly from the data by maximising the likelihood function (equation 4). The density of the matrix-variate *T* distribution is given by:

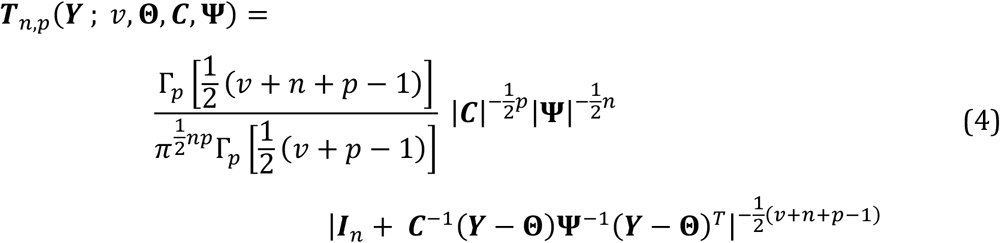

Where Γ_*p*_is the multivariate gamma function of dimension *p,* and ***I***_*n*_ an identity matrix of dimension *n* (Gupta and Nagar 2000; Iranmanesh et al. 2010). Because ***R*** has been integrated out, fitting a multivariate model of trait evolution using the marginal likelihood *T* (equation 4) alleviates the need to work directly with the estimate ***R̂***, and allows manipulating instead the simpler (typically diagonal – see above) matrix **Ψ**. Also, even though the equation (4) requires the computation of a determinant term involving the cross-product of a potentially large matrix, this can be achieved efficiently (see Supplementary material). Within this framework, fitting a model requires optimising equation (4) for the parameters controlling the phylogenetic variance-covariance matrix ***C***, and *μ*, one of the parameters defining the matrix **Ψ**.

### Regularised estimator of the variance-covariance matrix **R**

The estimation and storage of the variance covariance matrix ***R*** is not necessary for fitting multivariate models using equation (4). However, if one is interested in obtaining a regularised estimator of ***R***, this can be derived from its posterior distribution, which follows an inverse Wishart distribution (Iranmanesh et al. 2010; Thompson et al. 2020):

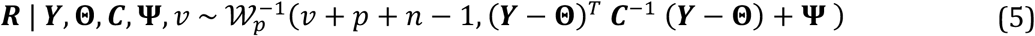

Given that the mean of an Inverse Wishart distribution of parameters *k*, **ψ** and dimension *p* is given by 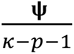 (when *κ* > *p* + 1; Gupta and Nagar 2000), the mean of the posterior distribution of ***R*** is given by:

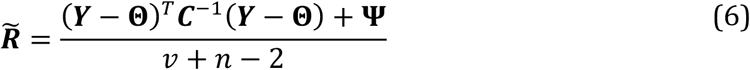

***R̃*** corresponds to the minimum mean square error estimator of ***R*** (MMSE) and is therefore a good regularised estimator candidate (Coluccia 2015). Note that under our assumption *v* = *p* + 1, the denominator in equation (6) scales with both *n* and *p,* as do several classical regularised estimators used in non-phylogenetic contexts (see for instance, Ledoit and Wolf 2004).

### Implementation

We implemented this approach for four widely-used models with high-dimensional versions: Brownian Motion (BM), Early Burst (EB), Pagel’s lambda, and Ornstein-Uhlenbeck (OU). Additionally, we extended the previously available high-dimensional models by implementing the Ornstein-Uhlenbeck process with multiple optima (OUM). In the OUM model, an optimum value of traits is estimated per selective regimes (i.e., group of lineages that are subject to a common selective pressure, defined *a priori* and reconstructed independently (Butler and King 2004)). Estimate of these *m* optima can be solved using equation (2) with the design matrix ***X*** replaced by a *n* by *m* weight matrix computed following Butler and King (2004) and Clavel et al. (2015). Because our formulation relies on the Kronecker structure – that is, a single ***C*** and ***R*** matrix are used to characterise the models – we assume that the same strength of selection α is acting independently on each trait in the OU models (i.e., the matrix ***A***, the multivariate counterpart to the parameter α (Clavel et al. 2015), is a diagonal scalar matrix), following Bastide et al. (2018) and Clavel et al. (2019). As with the other models, the marginal likelihood is given by equation (4), and the posterior mean of the variance-covariance matrix is given by equation (6).

We Bit the models by maximising the restricted likelihood (REML) to account for the degrees of freedom lost from fixing the root/mean for each trait, directly estimated from equation (2) (i.e., the fixed effect coefficients in regression models; Patterson and Thompson 1971; Gurka 2006). The REML provides an unbiased estimator of the parameters, improving the inference over the maximum likelihood (ML), especially when the sample size is small (Patterson and Thompson 1971) and/or in high-dimensional cases (Clavel et al. (2019) and results not shown). The REML is obtained by adding the terms |***X***^*T*^***C***^−1^***X***|^*p*^ and |***X***^*T*^***X***|^*p*^ to equation (4) with *n* corrected by the number of mean values *m* estimated by trait (i.e., the number of columns of the design matrix ***X***) (Harville 1974). For all the models considered here except the OUM, only one mean value is estimated by trait and thus, *m=1* and the term |***X***^*T*^***X***|^*p*^ is a constant that can be omitted when maximising the REML. For the OUM, because *m* is equal to the number of optima (i.e., the number of selective regimes) and the design matrix ***X*** depends on the parameter α controlling the strength of selection (Butler and King 2004), the term |***X***^*T*^***X***|^*p*^ cannot be omitted. The restricted log-likelihood is then proportional to:

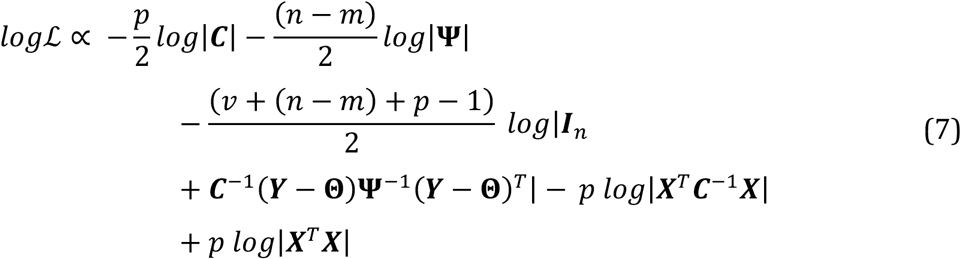

We maximize this log-likelihood function using a quasi-Newton method (by default the L-BFGS-B algorithm (Byrd et al. 1995) is used) implemented in the *optim()* function in R.

We implemented the Empirical Bayes approach in the *mvgls()* function (*method=’EmpBayes’*) from the R package ‘mvMORPH’ (Clavel et al. 2015). The proposed method allows incorporating extinct species, accounting for unknown measurement errors by jointly estimating a nuisance term, and computing confidence intervals using the Fisher information matrix (Supplementary material, Fig. S2). We provide a tutorial code for using the Empirical Bayes approach and the related functions in the Supplementary Material.

### Model selection using the Empirical Bayes approach

The marginal distribution of the Bayes model (equation 3) is a parametric distribution with (hyper)parameter **Ψ**, hence the framework of Information Criteria can be used (Konishi and Kitagawa 2008, section 9.1). We considered the Akaike Information Criterion (AIC, Akaike 1974), the Bayesian Information Criterion (BIC, Schwarz 1978), and the Extended Information Criterion (EIC, Ishiguro et al. 1997; Kitagawa and Konishi 2010) as options to select the proper (i.e., simulated) model of traits evolution. The AIC and BIC criteria are more widely used than the EIC, which uses a bootstrap resampling procedure to estimate the bias term and is therefore more intensive computationally. However, EIC is more accurate in general (non-asymptotic) conditions (Ishiguro et al. 1997; Kitagawa and Konishi 2010), and usually works better when the number of observations is small compared to the number of parameters (see Schwarz (1978); Konishi and Kitagawa (2008) for an extended review on all these criterions). We incorporated the calculation of the information criteria for the Empirical Bayes approach in the functions *AIC()*, *BIC()* and *EIC()* from ‘mvMORPH’. We also considered the bootstrap-based Likelihood Ratio Tests (LRT) (Williams 1970; Lewis et al. 2011) as an alternative to test the support for an evolutionary model relative to another one. While the well-known parametric LRT is restricted to nested models, bootstrapping techniques allow its application to non-nested model comparison (Lewis et al. 2011; Boettiger et al. 2012; Goolsby 2016). The bootstrap-based LRT relies on the comparison of a likelihood ratio, calculated from the observed data by *LR*_*obs*_ = −2 ∗ (log *likelihood null model* − log *likelihood alternative model*), against a null distribution of the test statistic *LR*_*boot*.*null*_ estimated by refitting to data simulated by bootstrap resampling (Lewis et al. 2011). The LRT is implemented in the *LRT()* function in ‘mvMORPH’. We follow Clavel and Morlon (2020) to perform the bootstrap for both EIC and LRT. This is achieved by sampling with replacement the normalised residuals of the fitted model, 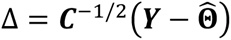, to obtain bootstrap samples 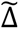. These samples are then used to generate new datasets, 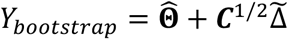 This bootstrap procedure enables us to generate random datasets with the same phylogenetic structure as the fitted model (whether null or alternative for the LRT) while retaining the same covariance structure between traits as in the empirical data. We performed this bootstrap procedures *n-times* to generate data under the null (or alternative) model for obtaining the distribution of the LRT statistic or for computing the EIC. Finally, for all the information criteria and the LRT, we evaluated the likelihood (equation 4) at the REML estimates to facilitate the comparison of models that differ by their design matrix ***X*** (i.e., different mean structure) (Verbyla 2019). For this, the likelihood for the REML estimates is re-calculated using the argument ‘*REML=FALSE’* in each function.

### Testing the performances of the Empirical Bayes approach through simulations

We used simulations to assess the performances of the proposed Empirical Bayes approach in terms of parameter estimation accuracy, model selection, and computational efficiency. We compared the performances to the penalized likelihood (PL) approaches that were shown to outcompete other methods dealing with high-dimensional data, and are the only ones explicitly providing an estimate of the variance-covariance matrix ***R*** (Clavel et al. 2019). In the Supplementary material, we also provide a comparison with the pairwise composite likelihood (PCL, Goolsby 2016), and with a naive implementation, where traits are assumed to be independent and the parameters are inferred by maximising the sum of the log likelihoods estimated for each trait independently.

For the PL approaches, we considered the archetypal ridge (PL-AR) and the quadratic ridge penalisations (PL-QR), using the *mvgls()* function from ‘mvMORPH’ (penalties options ‘RidgeArch’ and ‘RidgeAlt’ respectively) (Clavel et al. 2019). The penalisation target for both Ridge-type PL was chosen to match the scale matrix **Ψ** of the Inverse Wishart distribution in the Empirical Bayes approach. We used the ‘unitVariance’ and ‘Variance’ target option in *mvgls(),* for the scaled identity **Ψ**_***iden***_ and the diagonal **Ψ**_***diag***_respectively. We also used the ‘null’ target for the PL-QR because it showed the best performance among the rotation-invariant estimates in a previous study (see Clavel et al. 2019). We set the same optimisation bounds for all the approaches and we obtained the initial parameters for the numerical optimisation through a preliminary grid search.

We generated multivariate traits from classical models of trait evolution (equation 1) from simulated phylogenetic trees and covariance matrices ***R***. We simulated trees of size 100 (i.e., the number species *n* = 100) from a pure-birth process using the *pbtree()* function from the ‘phytools’ R package (Revell 2012, 2024), which we scaled to unit height. We generated positive definite covariance matrices ***R*** using two different strategies : i) using an ***UDU***^*T*^ (eigen) decomposition, where ***D*** is a diagonal matrix with values drawn from an exponential distribution with rate=0.01, and ***U*** are randomly oriented orthogonal eigenvectors matrices (Uyeda et al. 2015); and ii) sampling from a Wishart distribution, with *p* degrees of freedom and a scale matrix constructed using a separation strategy (e.g., Barnard et al. 2000) with a correlation matrix (i.e., matrix with off-diagonal entries set to 0.7 and diagonal entries to 1) scaled by a vector of variances sampled from a log-normal distribution (mean=3; standard deviation=0.8). While the first approach allows a better control on the distribution of the eigenvalues, it tends to generate datasets with low correlations between traits (Fig. S1). In contrast, sampling from the Wishart distribution allows a better control on the average level of correlation between traits, but with skewed eigenvalue distributions (Fig. S1). Using both strategies, we generate heteroscedastic datasets (i.e. heterogeneity of traits variances). Because of these properties, hereafter we refer to the datasets generated by the first strategy (eigen decomposition) as *weakly-correlated*, while those generated by the second one (from a Wishart distribution) as *highly-correlated*. We used the function *Posdef()* from ‘RPANDA’ (Morlon et al. 2016) package in R to generate the *weakly-correlated* covariance matrices, and custom codes for generating the *highly-correlated* covariance matrices.

We simulated traits following four evolutionary models: Brownian motion (BM), Ornstein-Uhlenbeck under different selection strengths (α = 0.34, 1.2 and 2.7) with one (OU) and multiple optima (OUM, 2 optima), Early-Burst (EB) with different decay rates (*r =* -0.34, -1.2 and -2.7), and Pagel’s lambda (λ = 0.2, 0.5 and 0.8), using the function *mvSIM()* from ‘mvMORPH’. For the OUM, we assigned the species to one of two selective regimes defined by a stochastic process with equal transition rates (rate = 0.3), using the function *sim.history()* from the ‘phytools’ package, ensuring that at least 35% of the species are represented in both of the two regimes. Next, we simulated traits with different means for each group (β_1_=-20 and β_2_=20). For all the other models, the mean was set to 0. We chose both, the selection strengths and the decay rates for OU and EB models respectively, for representing 2, 0.5 and 0.25 half-lives of the simulated phylogenetic trees. We evaluated the performances for high-dimensional datasets, with a number of traits ranging from twice, five, to ten times the number of species (i.e., *p* = 2*n* =200, *p* = 5*n* = 500, and *p* = 10*n* = 1000). We simulated 100 datasets for each combination of parameters. We provide the R script used to generate the simulations in a Zenodo repository (see *Data availability statement*).

We compared the similarity between the simulated matrix ***R*** and the regularised estimate of the covariance matrix (***R̃***), using the Kullback-Leibler (KL) divergence (equation (S1); Kullback and Leibler (1951)), and a quadratic loss (QL) function (equation (S2); Van Wieringen and Peeters (2016)). These two divergence measures differ in the statistical loss they capture: the KL divergence evaluates the underlying distribution of the traits from their covariance matrices and is affected by the distribution of their eigenvalues (note the determinant and trace terms in equation S1), while the QL function measures the squared distances between the matrices, therefore emphasising large distances from both the diagonal and off-diagonal elements (see equation S2).

For model selection, we evaluated the efficacy of the AIC, BIC and EIC, and the LRT, to select the simulated model of trait evolution as the right one. To reduce the number of comparisons, we performed them for only one parameter for each evolutionary model (Brownian Motion -BM-, OU and OUM with α = 2.7, and EB with *r =* -2.7). Note that BM is equivalent to Pagel’s lambda model when *λ* = 1. For the EIC, which requires a computationally demanding bootstrap procedure, we reduced the number of species to *n* = 50, the number of traits to *p* = 2*n*, and we considered only *weakly-correlated* traits datasets, expecting a similar performance for *highly-correlated* traits given the results for the other information criteria (see *Results* section). We performed 100,000 bootstrap replicates, in order to obtain a stable and accurate estimation of the criterion (i.e., a standard error of the bias term below one unit). For the information criteria, we evaluated the consistency of the model selection by counting the number of times the true model is selected by the criterion, while for the LRT, we measured the number of times the true model is not rejected (p-value >= 0.05, when it is used as a null hypothesis; or p-value < 0.05, when used as alternative). Following Boettiger et al. (2012), we define the null model as the less parameterised one, unless the models have the same complexity (number of parameters) (Table 1).

**Table 1.**
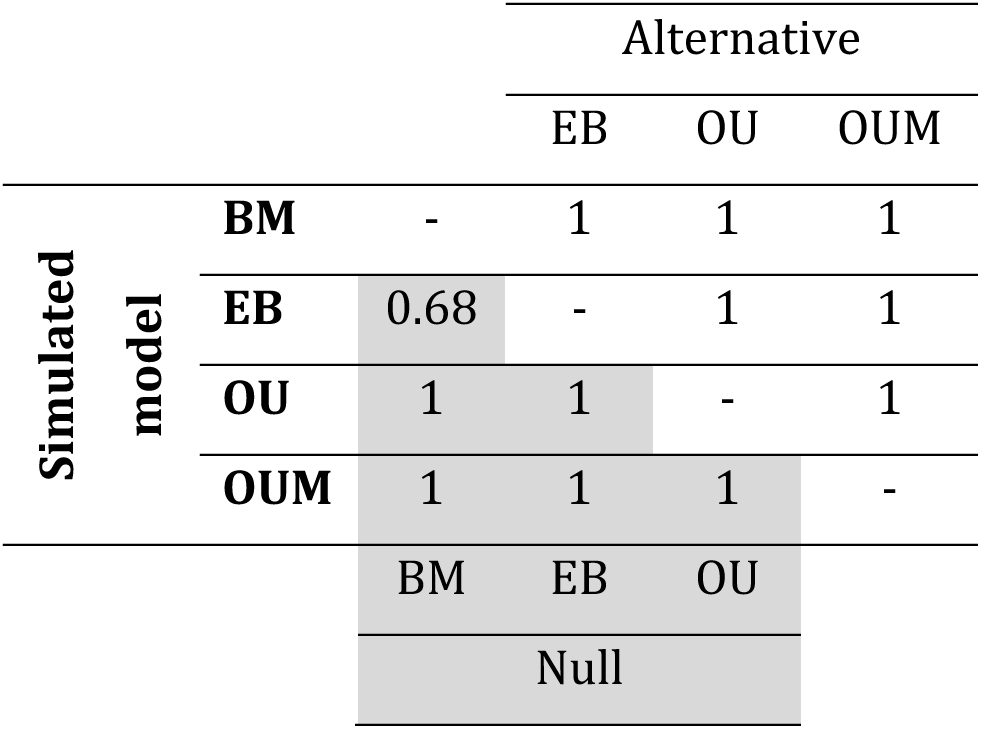
Proportion of times that the generating model is supported across 100 simulations according to the Likelihood Ratio Test (LRT). Support is evaluated by not rejecting the null hypothesis when it corresponds to the simulated model and by rejecting it when the true model is used as the alternative hypothesis. The threshold for significance level is α=0.05. The grey area (lower triangular) indicates the cases when the simulated model is used as the alternative hypothesis in the LRT, while the non-coloured area (upper triangular) shows the results when the model is used as the null hypothesis. Because Brownian motion (BM) is the less parameterised model, it is always used as the null hypothesis and Ornstein-Uhlenbeck with multiple optima (OUM) as the alternative. EB and OU refer to Early-Burst and Ornstein-Uhlenbeck models, respectively. For the simulations *n*=50 and *p*=100.

### Computational performances

We evaluated the computational performance by fitting the Pagel’s lambda model (λ=0.9) over 50 replicates to a 50-species tree, and an increasing number of traits (*p=100, p=200, p=1000, p=2000, p = 4000).* To save computational resources, we only considered simulations with a *weakly-correlated* covariance matrix ***R***. We benchmark the Empirical Bayes approach to the two PL methods (PL-AR, PL-QR) with the **Ψ**_***iden***_ target, all implemented in the *mvgls()* function. For a fair comparison, we fixed the same initial values for the optimisation for all the approaches. For the PL with archetypal ridge (PL-AR), we used the leave-one-out cross-validation (LOOCV) strategy used to estimate the regularisation parameter, as well as a faster implementation based on rank-one update of the LOOCV proposed by Hoffbeck and Landgrebe (1996) (‘*H&L’* method in *mvgls()*; see details in Clavel et al. (2019)). For the PL-AR using the LOOCV and PL with quadratic ridge (PL-QR) we only evaluated the computing time for *p=100, p=200 and p=1000*, due to the computational burden associated with larger dimensions. Because the estimation of the regularised covariance matrix ***R̃*** for the Empirical Bayes is not required in the model fitting process but it is estimated by default in *mvgls()*, we assessed both cases (with and without its explicit computation). We used the ‘bench’ package (Hester and Vaughan 2025) to evaluate the total time and total memory allocated by R when fitting a model. We ran the *mark()* function repeatedly on each approach and for each simulated dataset, executing it on a single processor each time. We ran the simulations at the Centre de Calcul de l’IN2P3 (CC-IN2P3/CNRS, https://cc.in2p3.fr), on a Red Hat Enterprise Linux release 9.7 (Plow) system with an architecture x86-64, and R version 3.4.4.

### Applying the Empirical Bayes approach to empirical data: The injluence of diet in the evolution of terrestrial mammals’ lower jaws

We applied the newly developed models to test hypotheses related to adaptive convergence in mammalian lower jaw shape in relation to dietary regimes, and constraints such as those imposed by extended suckling during early development in metatherians *versus* eutherians. We used the dataset published by Fabre et al. (2021), consisting of 114 Procrustes aligned 3D landmarks and sliding semi-landmarks (*p =342*) describing the lower jaw shape. To avoid overparameterisation problems and lack of statistical power, we used only the diet categories from Fabre et al. (2021) that are represented by at least 10 species in each of both Metatheria (marsupials and their stem relatives) and Eutheria (placentals and their stem relatives). We focused on two broad diet categories: animal-based (carnivorous) and plant-based diets (herbivorous). The *Carnivorous* regime is composed of ‘Insectivorous’ and ‘Carnivorous’ species while the *Herbivorous* regime groups together the ‘Browser’, ‘Grazer’ and ‘Mixed feeder’ categories from the original dataset (Fabre et al. 2021). The final dataset consists of 95 species (74 extant and 21 extinct): 43 carnivorous and 52 herbivorous lineages, including 8 and 13 fossils lineages respectively. We assessed the support for two hypotheses on the diet regimes favouring morphological differences, between: 1) carnivorous and herbivorous lineages (2 optima, OUM2); and 2) carnivorous and herbivorous lineages, but with different optima for Eutheria and Metatheria clades (4 optima, OUM4_cl_). We also assessed the support for one hypothesis on the development mode constraining the jaw morphology and leading to 3) two different optima, one for Eutheria and another one for Metatheria (OUM2_dv_). Finally, we evaluated the Bit of a single optimum OU (i.e., an OU model with no differences in shapes between clades, diet regimes, or other ecologies), an Early-Burst (EB) model representing declining rates of trait evolution as species diversification proceeds, and a Brownian motion (BM) model. We used the same phylogenetic tree as Fabre et al. (2021) trimmed to our final dataset. Although the Procrustes superimposition removes scale effects, we note that allometric trends may still exist, meaning that changes in the lower jaw shape may be related to changes in size.

We based the hypotheses for the OUM models on the joint reconstruction of the ancestral diet states under a continuous time Markov chain (Mk model, (Lewis 2001; Revell 2025)). We performed the reconstruction with the function *ace()* from ‘ape’ R package (Paradis and Schliep 2019), using a model with symmetrical transition rates across states (SYM), which showed the best Bit compared to the models with unequal and asymmetrical rates (ARD) or with equal rates model (ER). We fitted the models using the rotation invariant scaled-identity target matrix to fulfil the requirements of geometric morphometrics data (Rohlf 1999), allowing the estimation of measurement errors and confidence intervals (‘*error=TRUE’*, ‘*FCI=TRUE’* respectively, in *mvgls()*). We computed the AIC, BIC and EIC to assess the model Bit to the data, using the likelihood (rather than the REML, as explained above) evaluated at the REML estimates (‘REML=FALSE’ in *AIC(), BIC(),* and *EIC()* functions in ‘mvMORPH’), and performing 3000 bootstrap resampling. We also compared the support of the OU and OUM models using the likelihood ratio test (LRT). Likewise, for the information criteria, we used the likelihood evaluated at the REML estimates and performed 3000 bootstrap resampling to generate the null distribution. To visualize the estimated optima, we mapped the inferred coordinates onto one of the mesh of the 3D scans from Fabre et al. (2021) using the function *tps3d()* from ‘Morpho’ package in R (Schlager 2017), and then we generated a 3D surface reconstruction using *shade3d()* from ‘rgl’ package (Murdoch and Adler 2025). All the plots were generated using multiple functions from the ‘ggplot2’ package (Wickham 2016).

## Results

### Parameter inference

The Empirical Bayes approach accurately inferred the evolutionary parameters for all the models (i.e. Pagel’s lambda, *r* for EB, or α for OU), regardless of the structure of the covariance matrices simulated, the target matrix, and the number of traits *p* (Fig. 1, S3, S4). The Penalized Likelihood (PL) showed similar performance (Fig. 1, S3, S4), while the pairwise composite likelihood (PCL) showed a higher variance, and the naive inference assuming trait independency showed both higher variance and low accuracy in parameter estimates (Fig. S3-S4).

**Figure 1.**
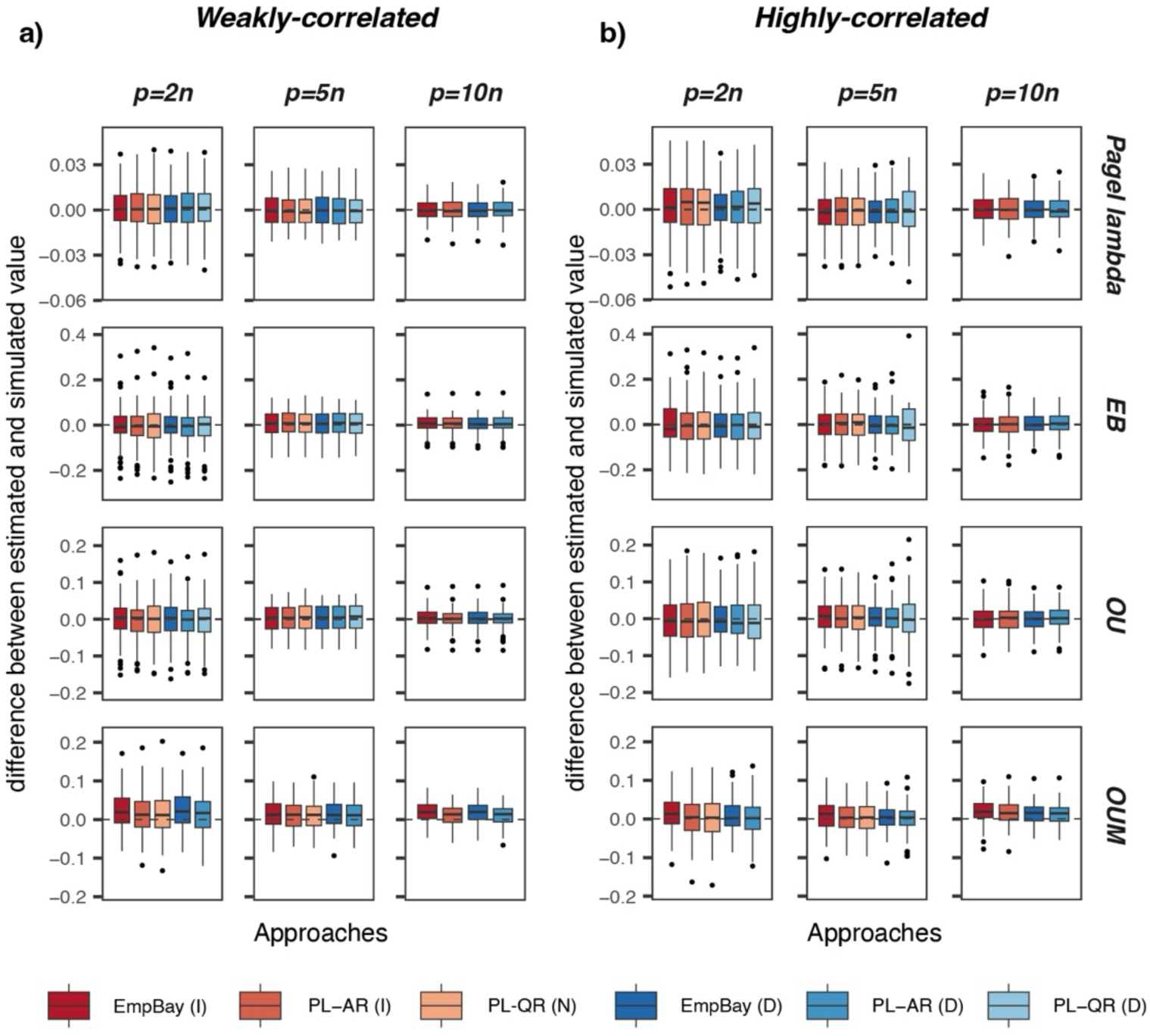
Difference between the estimated and the simulated parameter for four evolutionary models: Pagel’s lambda (λ= 0.5), Early Burst (EB, r=-1.2), Ornstein-Uhlenbeck (α=1.2) with one (OU) and multiple optima (OUM), for traits (**a**) *weakly-correlated* and (**b**) *highly-correlated*. All the approaches estimate the parameters accurately (values around zero, indicated by a dashed line), irrespective of the ratio between the number of traits and the number of species (*n*=100), across 100 simulations. Approaches: EmpBay = Empirical Bayes; PL-AR: Penalized likelihood – Archetypal Ridge; PL-QR: Penalized likelihood – Quadratic Ridge. Target matrices: scaled-identity (I), null (N) and diagonal (D) matrix. The PL-QR analyses were not run under some conditions because of their computational cost. Results for the other parameter values are reported in Figures S3 and S4.

For Pagel’s lambda, OU and EB models, the variance in the parameter estimates decreased with an increasing ratio between the number of traits and the number of species (*p/n*) (Fig. 1). For the OUM model, the accuracy increased with increasing α values (strength of selection) with a slight underestimation at the lowest value (Fig. 1, Fig. S3-S4).

### Estimation of the variance-covariance matrix **R**

The accuracy of the (regularised) variance-covariance estimates depends on the data structure (*weakly-correlated* or *highly-correlated* traits), the choice of target matrix, and the choice of similarity measure (i.e. the Kullback-Leibler (KL) divergence or the Quadratic Loss (QL)) (Fig. 2, S5-S8). For *weakly-correlated* traits simulations, the Empirical Bayes and the PL estimators showed comparable KL distances to the true (simulated) variance-covariance matrices ***R*** with either target, except for the PL with quadratic ridge (PL-QR) with null target, which showed a higher divergence (Fig. 2a, S5). In general, the use of one or the other target matrix resulted in noticeable differences only in the datasets with the highest *p/n* ratio, where the approaches using the scalar identity matrix showed a slightly lower divergence on average (Fig. 2a, S5). In terms of QL, the approaches using the scalar identity target matrix also showed a lower loss compared to those using the diagonal target, except for the PL with quadratic ridge (PL-QR), which had the lowest loss when using the diagonal target, outperforming all the other approaches when it was possible to use it (i.e., when computing time was not prohibitive, Fig. S7). The Empirical Bayes estimates showed a lower QL loss at the lowest *p/n* ratio evaluated, but this difference vanished as *p/n* ratio increased, with a similar performance to the PL-AR (Fig. S7). For *highly-correlated* traits, we found markedly different results between the approaches depending on which target matrix was used, with both the KL divergence and QL distance measures (Fig. 2b, S6, S8). With KL, divergences were overall lower with the diagonal matrix target than with the scalar identity target matrix (Fig. 2b, S6), while the opposite was true with QL (Fig. S8).

**Figure 2.**
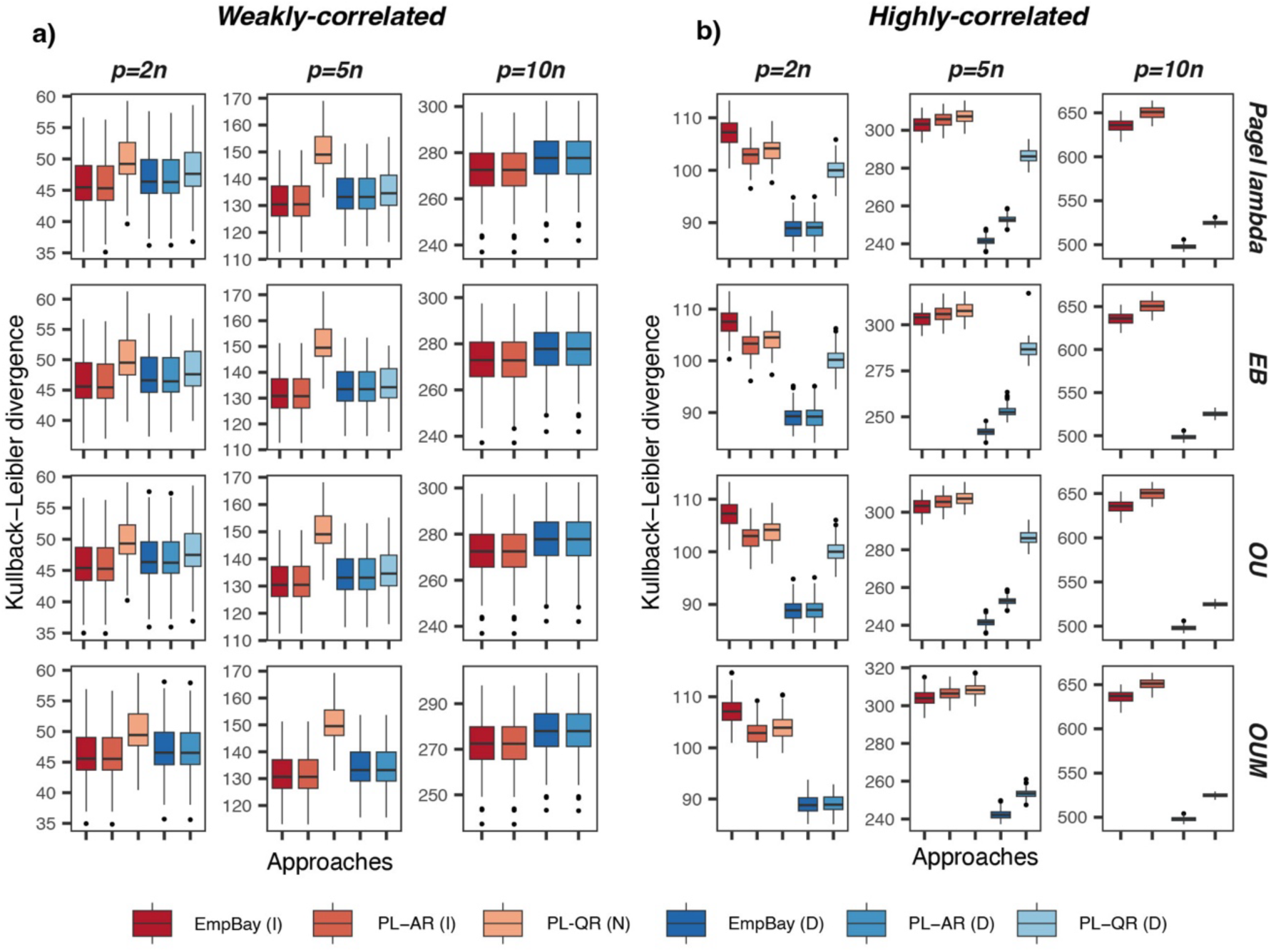
Kullback-Leibler divergence between the estimated and simulated traits covariance matrix under four evolutionary models: Pagel’s lambda (λ= 0.5), Early Burst (EB, r=-1.2), Ornstein-Uhlenbeck (α=1.2) with one (OU) and multiple optima (OUM), for traits (**a**) *weakly-correlated* and (**b**) *highly-correlated*. The lower the divergence, the more similar are the matrices. Boxplots summarize the results for 100 simulations with three different ratios between the number of species (*n*=100) and the number of traits ranging from twice (*p=2n*) to ten times (*p=10n*). Approaches: EmpBay = Empirical Bayes; PL-AR: Penalized likelihood – Archetypal Ridge; PL-QR: Penalized likelihood – Quadratic Ridge. Target matrices: scaled-identity (I), null (N) and diagonal (D) matrix. The PL-QR analyses were not run for the simulations with *p=10n* because of their computational cost. Results for the other parameter values are reported in Figures S5 and S6, and results with the Quadratic Loss are reported in Figures S7 and S8.

### Model selection

Across all our model comparisons, we found that the EIC was the most reliable criterion (total misclassification score of 8%, compared to 11.5% for the BIC and 14% for the AIC). With the AIC and BIC criteria, we correctly selected the right model, except for datasets simulated under Brownian motion (BM). In this situation, the Ornstein-Uhlenbeck (OU) was favoured over BM around half of the time (Fig. 3, Fig. S9). The EIC correctly selected the right model most of the times, including the BM model (92/100), although it was less efficient than AIC or BIC for traits simulated with the Early-Burst (EB) model, where the OU with multiple optima (OUM) was incorrectly favoured around 20% of the time (Fig. 3). The likelihood ratio test (LRT) enabled differentiating the correct model (i.e., non-rejection of the null hypothesis when the true model is used as the null, or rejection when it is used as the alternative model) for nearly all comparisons, except for traits simulated under an EB process, where the EB model was favoured over BM only 68% of the time (Table 1).

**Figure 3.**
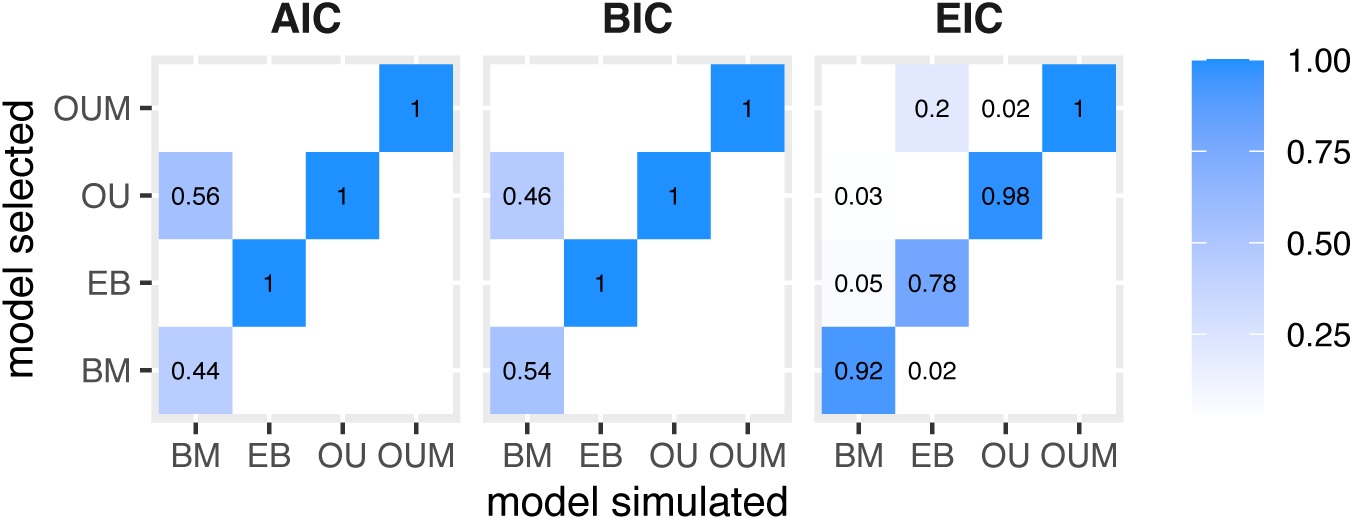
Proportion of times the simulated model was correctly selected (lowest information criterion value) using Akaike (AIC), Bayesian (BIC) and the Extended (EIC) Information criteria, across 100 simulations, when the number of traits is twice the number of lineages (*n*=100; *p*=200) and the traits are *weakly-correlated*. Models: Brownian Motion (BM), Early Burst (EB), Ornstein-Uhlenbeck with one (OU) and multiple optima (OUM). Results for the other dimensions and for traits *highly-correlated* are reported in Figure S9.

### Computational efjiciency

The Empirical Bayes approach reduced the computing time for fitting the multivariate models compared to the PL approaches. For the largest dimensions (*n = 50; p = 4000*) the Empirical Bayes approach was up to approximately ten times faster than the fastest PL implementation (i.e., using Hoffbeck and Landgrebe (1996) fast LOOCV computation). This difference was even higher for the PL using the standard LOOCV, where the Empirical Bayes approach was up to three orders of magnitude faster than the PL-QR when the number of traits is twenty times the number of lineages (*p = 1000,* Fig. 4). Since it is not required for the inference, the estimation of the matrix ***R̃*** is optional in our current implementation. When it is not estimated, the computing time is reduced by up to two orders of magnitude (approximately 200 times) compared to the fastest implementation of the PL at the largest evaluated dimensions (Fig. 4a, Table S2).

**Figure 4.**
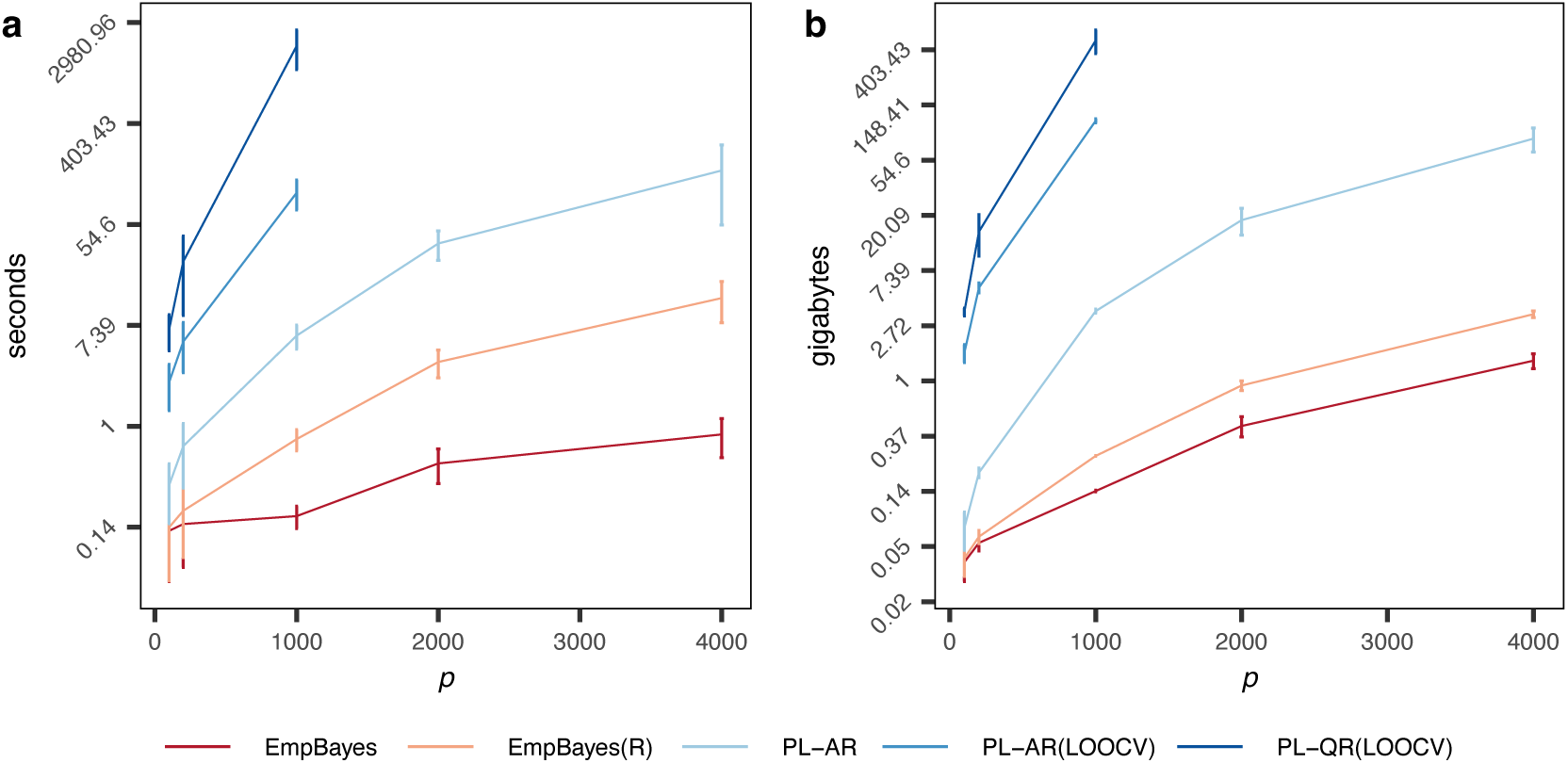
Computational performances in terms of **a)** total execution time (seconds) and **b)** memory allocated by R (gigabytes - GB), when fitting Pagel’s lambda model (λ=0.9). Both plots use a logarithmic scale while preserving the absolute values in the labels (all the values in Table S2). The Empirical Bayes approach has the best performances in terms of both computing time and memory use, with greatest improvement at largest (*p*) dimensions. The plots show the results across 50 simulations, when *n*=50 and the number of variables increases from *p=n* (50) to *p=80n* (4000). Approaches: EmpBayes: Empirical Bayes approach; EmpBayes(R): EmpBayes with the explicit computation of the matrix ***R̃***; PL-AR: Penalized likelihood with archetypal ridge penalty estimated using a fast LOOCV implementation; PL-AR (LOOCV): PL-AR using the regular leave-one-out cross-validation method; PL-QR (LOOCV): PL with quadratic ridge penalty estimated using the regular LOOCV (this is the only possible approach available for the PL-QR).

The Empirical Bayes approach was also more efficient than the PL approach in terms of memory use. The total amount of memory used by each approach (i.e., the sum of the memory used by all the expressions called in the functions) is presented in Figure (4b). At the largest dimensions (*n = 50; p = 4000*), the Empirical Bayes approach required up to 20 times less memory than the PL-AR when computing the matrix ***R̃***, and up to 50 times less memory when not computing it. For instance, in our simulation with p=4000, the Empirical Bayes approach required around 3.3 GB on average when computing the matrix ***R̃*** explicitly, while the PL-AR with the fast LOOCV used around 80 GB. This difference is even greater between the Empirical Bayes and the PL approaches when a regular leave-one-out cross-validation procedure is used (PL-AR and PL-QR LOOCV; Fig. 4b, Table S2). For p=1000, the difference is up to three orders of magnitude (Fig. 4b, Table S2). As the computing time and memory usage do not scale linearly with the number of traits, the efficiency of the Empirical Bayes is particularly noticeable from dimensions larger than p=200, with the relative improvement increasing steadily (Fig. 4).

### The injluence of diet on the evolution of terrestrial mammals’ jaws

The reconstruction of the ancestral states for the diet regimes (Fig. 5a) suggests an ancestral carnivorous state and repeated evolution toward herbivory (up to 7 times in the studied sample). The evolution of jaw morphology across mammals was best supported by an Ornstein-Uhlenbeck process with two optima associated with diet, according to two of three model selection criteria we used: while the BIC favoured the simpler OU with a single optimum (OU), the AIC and the more trustable EIC favoured the OUM2 scenario (carnivorous/herbivorous, regardless of whether the species are metatherian or eutherian) (Table 2). Furthermore, hypothesis testing with the LRT rejected the OU with a single optimum when compared to the OUM2 (LRT_Null:OU-Alt:OUM2_ = 1104.43, p-value=0.003), but not when compared to the OUM2_dv_ (i.e., evaluating selection leading to different optima for Metatheria and Eutheria, LRT_Null:OU-Alt:OUM2dv_ = -112.0885, p-value=0.885), or the OUM4_cl_ (i.e., different morphologies between marsupials and placentals in addition to differences associated with diet; LRT_Null:OU-Alt:OUM4cl_ = 1416.269, p-value=0.313). Likewise, the OUM2_dv_ did not significantly improve model Bit when compared to the OUM2 (LRT_Null:OUM2-Alt:OUM2dv_ = -1216.518, p-value=0.31), nor did the OUM4_cl_ (LRT_Null:OUM2-Alt:OUM4cl_ = 311.8387, p-value=0.803) (Fig. 5b). While the LRT test rejected the OU model in favour of the OUM2, we note that the observed LR statistic did not fall within the 95% of either distribution but rather between them (Fig. 5b). This suggests that, although a single optimum OU is unlikely, the OUM2 may actually be slightly underfitting the data (e.g., some lineages might have their own optima). The OUM2 describes a model with a low selection strength (α≅0.0048, about 1.1 half-lives elapsing over the 168 Ma of mammalian history we considered, Table 2) leading to one optimum for carnivorous, and one for herbivorous, which differ mainly in the corpus depth (1 in Fig. 5c) and the shape of the posterior region of the mandible (2 in Fig. 5c). The herbivorous lineages tend to have a longer and deeper corpus (i.e., anterior region, 1) than carnivorous lineages. Also, the herbivorous optimum shows a taller ramus (i.e., posterior region, 2) with a larger masseteric fossa (6), narrower sigmoid notch (7), thinner coronoid process (3) and a less prominent angular process (5) than the carnivorous. Additionally, in herbivores the condylar process (4) is oriented dorsally, whereas in carnivorous lineages, the condylar process is oriented more caudally.

**Figure 5.**
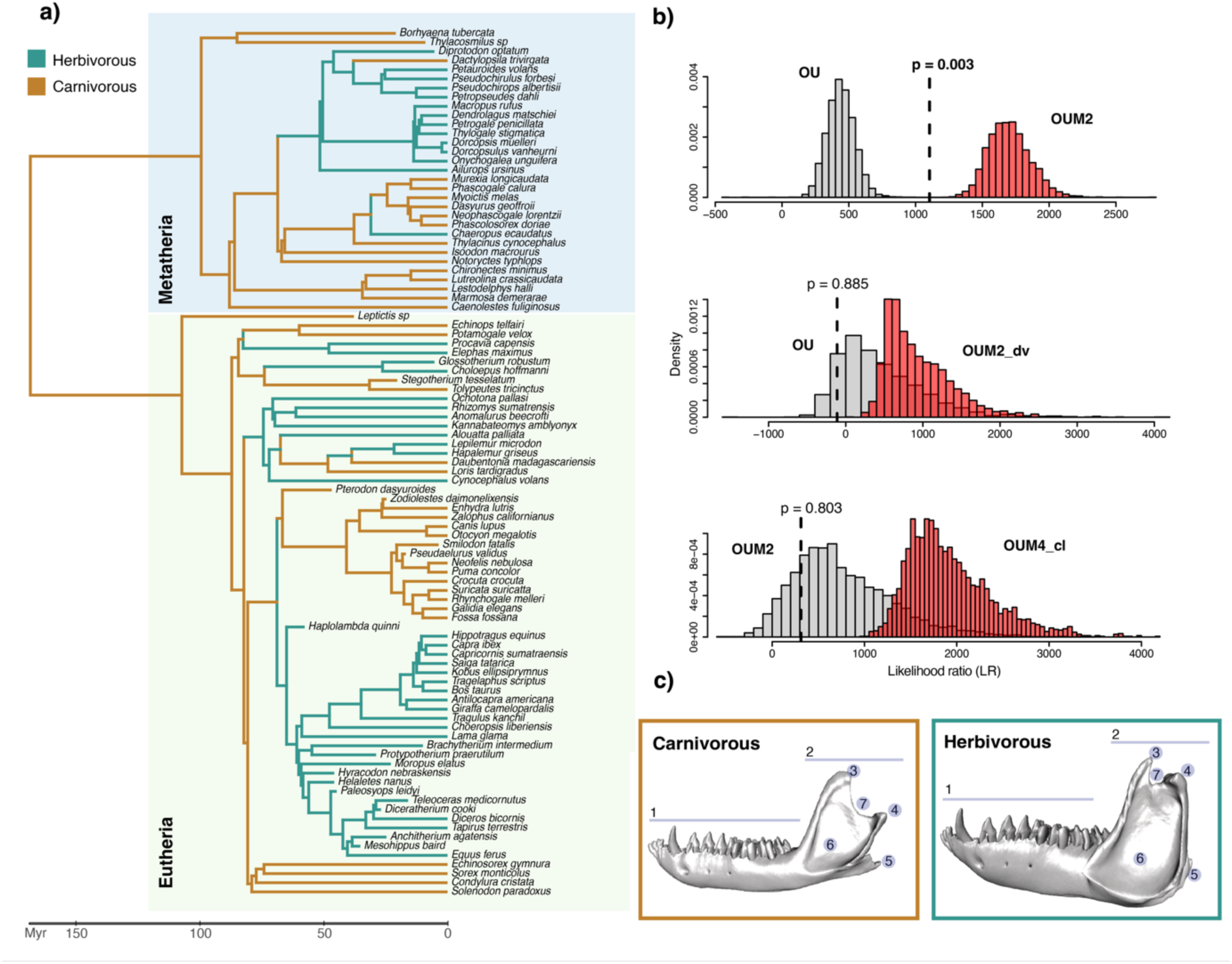
The adaptive convergent evolution of the jaw in herbivorous and carnivorous mammals. **a**) Phylogenetic tree for the 95 mammal species (Eutheria and Metatheria) analysed in this study. The most supported model, OUM2, is one which models the adaptation towards two distinct optima corresponding to the two diets. **b**) Results of the pairwise comparisons using the likelihood ratio test (LRT) for the OU models. The histograms show the distribution of the likelihood ratio obtained when the data were generated under each model (indicated next to each distribution). The dotted line indicates the observed LR statistic on the empirical dataset. Support for rejecting the null hypothesis (bars in grey) is indicated by the p-value. The LRT rejects the OU model in favour of the OUM2 (note the p-value and the location of the observed LR statistic relative to each distribution; top panel), although the observed LR falls in the lower tail of the distribution (red bars). **c**) Reconstruction of the optimum estimated for carnivores and herbivores under the OUM2 model. The main differences are located on the corpus depth (1), the positions of the coronoid (3), condylar (4) and angular (5) processes in the ramus of the jaw (2), and the extension of the masseteric fossa (6) and the sigmoid notch (7).

**Table 2.** Delta AIC, BIC and EIC for the models Bit on the mammals’ jaw dataset. The delta shows the difference between criteria value for each model and that of the model with the lowest value. The standard error (SE) around the estimated EIC value is also reported. The alpha parameter (selection strength) for the OU models is shown, with the confidence intervals within parenthesis. The AIC and EIC favoured the OUM model with two optima (herbivorous vs carnivorous), while the BIC favoured the OU with a single optimum. Note that the estimated confidence interval for alpha in the OUM4_cl_ produces a lower bound that is negative (-0.0013) replaced by zero. This numerical artefact has no biological significance but instead indicates that the uncertainty around the estimated parameter includes zero and cannot therefore be distinguished from it. BM: Brownian motion; OU: Ornstein-Uhlenbeck with one optimum; EB: Early-Burst; OUM: Ornstein-Uhlenbeck with distinct optima for herbivores and carnivores (OUM2), Metatheria and Eutheria (OUM2_dv_), and for herbivorous metatherian and eutherian as well as carnivorous metatherian and eutherian (OUM4_cl_).

| Model | $\Delta AIC$ | $\Delta BIC$ | $\Delta EIC$ | SE EIC | $\alpha$ |
| --- | --- | --- | --- | --- | --- |
| *BM* | -682.810 | -259.826 | -781.634 | 22.383 | - |
| *EB* | -684.806 | -264.376 | -877.73 | 22.788 | - |
| *OU* | -420.430 | **0.000** | -682.4 | 23.025 | 0.0043 (0.0032, 0.0055) |
| *OUM2* | **0.000** | -452.995 | **0.000** | 21.687 | 0.0047 (0.0035, 0.0058) |
| *OUM2<sub>dv</sub>* | -1216.518 | -1669.514 | -1474.543 | 22.888 | 0 |
| *OUM4<sub>cl</sub>* | -1056.453 | -3256.301 | -963.417 | 21.013 | 0.0008 (0, 0.0030) |

## Discussion

Here, we presented a new Empirical Bayes approach for modelling high-dimensional multivariate traits that avoids the explicit computation and storage of the ill-conditioned variance-covariance matrix. We showed that this approach accurately estimates evolutionary parameters for different models regardless of the level of correlation between the traits. We also showed that it is possible to inexpensively obtain a regularised estimate of the evolutionary covariance matrix between the traits, which outcompetes previously-developed regularised estimates based on penalised likelihood, in particular at large *p/n* ratios. The advantage of the Empirical Bayes method developed here (in comparison to the penalized-likelihood approaches) comes mostly from its computational efficiency, using up to 20 times less memory and being at least 10 times faster, which allow the extension to more complex models, as illustrated here with the multiple optima OU model, and the use of robust bootstrap or simulation-based procedures for testing model Bit.

### Parameter inference: An accurate and efjicient approach

Our results confirm the previously reported effectiveness of regularisation approaches for estimating and comparing evolutionary models with high-dimensional datasets (Clavel et al. 2019). By directly integrating the covariance between all the traits in the models, the Empirical Bayes approach – as well as the other methods based on regularisations of the covariance matrices – outperformed other phylogenetic comparative methods (PCMs) that only partially account for the correlations (i.e., PCL, Goolsby (2016)) or ignore them (naive implementation).

The main differences in performance among the various regularisation approaches come from the estimates of the covariance matrix ***R̂***. The Empirical Bayes and the PL with archetypal ridge (PL-AR) estimates show comparable performances, which can be explained by the similarity in the shrinkage performed by both approaches. Rewriting the Empirical Bayes estimator of the covariance matrix (equation 6) as ***R̃*** = ***R̂***(1 − *τ*) + *τ****J***, where 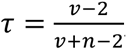, and 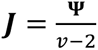, it is easier to see that it corresponds to a linear combination of the of the Inverse Wishart prior mean ***J***, with the maximum likelihood estimate for ***R***. This is similar to the regularisation performed by the PL-AR (see equation 3 in Clavel et al. (2019)). By contrast, the PL with quadratic ridge (PL-QR) performs a non-linear shrinkage of ***R̂***, changing its eigenvalues using a quadratic expression, thus offering more Blexibility and potentially superior performances than approaches based on linear shrinkages. When the distribution of the eigenvalues is not strongly skewed, shrinking them to the mean value (as done under linear shrinkage) brings them closer to their true values. Conversely, when the eigenvalue distribution is highly skewed (as in our simulation of *highly-correlated* traits), linear shrinkage over-shrinks the eigenvalues, bringing them far from their true values. Accordingly, our results suggest that the quadratic shrinkage performs better than our approach, but only for *highly-correlated* traits when similarity is measured using quadratic loss (QL) rather than Kullback-Leibler (KL) divergences. Under this extreme, skewed eigenvalue distribution, the regularisation applied by the Empirical Bayes approach probably shrinks the eigenvalues more heavily than the other approaches, reducing the characteristic dispersion of eigenvalues in high-dimensional settings (i.e. overestimation of the largest eigenvalues and underestimation of the smallest ones, see for instance Ledoit and Wolf (2020, 2021)). This severe regularisation generates a more similar eigenvalue structure, but also translates into larger QL distances than PL-QR, since ***R̃*** is pushed more severely towards the diagonal structure of the target matrices (i.e., no covariance among the traits). The effect of these large distances on the biological interpretation remains to be evaluated in future studies. Preferring a regularised estimator with a lower KL divergence or QL distances depends on the goal of the study (e.g., see Ledoit and Wolf 2021). For example, a similar eigenvalue structure may yield robust inference when ***R̃*** is used in multivariate analyses (e.g., see Clavel and Morlon (2020) for the linear shrinkage case with PL-AR). However, large distances due to strong regularisations (under the target matrix structure used here) may be undesired for studies aiming to describe patterns of covariance among the traits. In the current context of trait evolution, the proposed Empirical Bayes estimates offer the best compromise between computational feasibility and accuracy, since it is applicable in very high-dimensional settings where the PL-QR is not, and has performances which guarantee robust inference in subsequent multivariate analyses.

Our results underscore the importance of the target matrix used if the traits are highly correlated, which is expected in the case of high-dimensional datasets. For phenotypic data lacking a natural orientation in space, such as in geometric morphometric, the results of the model Bit should not depend on any arbitrary rotations of the data (Zelditch 2012; Lancewicki and Aladjem 2014; Adams and Collyer 2018). In these situations, the rotation-invariant identity matrix should be used as the target. Otherwise, the diagonal target matrix is appealing because it enables modelling different variances for each trait, potentially resulting in a better approximation to the true variance-covariance matrix. We found that using the diagonal target indeed provides an eigenvalue distribution closer to the true one, as shown by our results with the KL divergences, but potentially greater differences regarding the true values, as we found using the QL distances.

One of the most remarkable advantages of using the Empirical Bayes method (compared to other regularisation techniques) is the considerable reduction in computing time and memory requirements. This is achieved by marginalising the variance-covariance matrix ***R*** analytically, and inferring directly the *μ* parameter - controlling the level of regularisation-from the data without relying on computationally intensive cross-validation techniques. The approach circumvents the explicit and costly estimation, inversion and storage of the matrix ***R***. These operations were repeated multiple times by the cross-validation techniques used by the PL, resulting in approaches that were often unusable in practice (see Table S2). The gain in computational efficiency is particularly striking when the number of traits increases, making the Empirical Bayes approach amenable to large datasets (>4000 variables). This opens the possibility to model trait evolution from very-high dimensional comparative datasets such as those recently published in geometric morphometrics (e.g., Felice et al. 2019; Bardua et al. 2021; Goswami et al. 2022; Goswami and Clavel 2025), or to study other high-throughput quantitative traits that are typically analysed separately (e.g., Bedford and Hartl 2009; Chen et al. 2019; El Taher et al. 2020). A more efficient approach also opens the possibility to expand the PCMs to more complex models. These include models with a more complex mean structure, such as the OU model with multiple optima (OUM) implemented here, and other models considering the effects of climate (Clavel and Morlon 2017) or biotic interactions (Drury et al. 2016), which could be implemented in the future. Other regularisation approaches, such as PL, may be applicable to these models, but not for very large datasets. Indeed, the most efficient implementations of PL is based on computational tricks that can only be applied when a single ancestral state or optimum is estimated per trait (see Appendix 3 in Clavel et al. (2019)), restricting their applicability to simple models. For example, fitting the OUM to the mammalian jaws dataset (*n*=95, *p*=342) using PL-AR (LOOCV) was more than 60 times slower and required 155 times more memory than with the Empirical Bayes approach. Even for this relatively small dataset, applying some of the iterative procedures required for model comparison or statistical analysis in high-dimensional data may be prohibitive. We showed that the Empirical Bayes approach renders the use of the Extended Information Criterion (EIC) and the Likelihood Ratio Test (LRT) feasible, both of which are based on bootstrap resampling techniques. Similarly, the Empirical Bayes approach may benefit other multivariate analyses, such as MANOVAs and MANCOVAs, which are widely used for testing evolutionary hypotheses (e.g., Langerhans and DeWitt 2004; Da Silva et al. 2018; Kennedy et al. 2020; Goswami et al. 2022), but whose current regularised versions rely on costly permutation procedures (Clavel and Morlon 2020).

### The challenge of model selection

One of the current challenges for the study of phenotypic evolution from high-dimensional data is how to perform model comparison. The efficiency of the AIC and consistency of the BIC have been proved under asymptotic conditions, where *n* tends to infinity and *p* is fixed (Konishi and Kitagawa 2008; Ding et al. 2018). In our simulations, these two criteria were not able to select the Brownian Motion (BM) model when it was the generating process. This is not surprising given that BM is nested in OU (when the parameter α= 0, OU reduces to BM) (Hansen 1997; Butler and King 2004), and is consistent with previous findings where BM was often misidentified for an Ornstein-Uhlenbeck (OU) with weak support (Bartoszek 2016; Cooper et al. 2016; Adams and Collyer 2018; Clavel et al. 2019). On the other hand, the EIC has been shown to perform better across a broad range of conditions, including non-asymptotic ones (Ishiguro et al. 1997; Yafune et al. 2005; Kitagawa and Konishi 2010). For instance, compared to the AIC, it performs a higher (third-order) correction to the bias in approximating the expected likelihood from a finite (smaller) (Ishiguro et al. 1997; Kitagawa and Konishi 2010). Using the EIC In previous (PL) approaches was prohibitive because of the high computational cost associated to bootstrapping, but is now feasible with the Empirical Bayes approach. In our simulations the EIC showed the best performances for identifying the generating models, with the lowest performance for the Early-Burst (EB) model, confounded for an OUM model 20% of the time. If different regimes in the OUM arise early, divergences in trait pulled to different optima will also arise early, generating patterns that might resemble the early trait divergences of the EB model. Indeed, the EB and the OU with multiple optima (OUM) are both commonly used to model adaptive trait divergence into different ecological niches (Butler and King 2004; Harmon et al. 2010). This could explain incongruent results in some simulations for non-consistent criteria (i.e., which are not expected to select the correct model with probability equal to 1), such as the AIC and the EIC. Another approach made possible by the computational efficiency of the Empirical Bayes is the bootstrap-based likelihood ratio test (LRT), which was successfully used in previous PCMs for which devising an information criterion or evaluating the support of alternative models was difficult (Boettiger et al. 2012; Goolsby 2016). In our simulations, we found that the bootstrap-based LRT was useful for selecting the true model. However, LRTs are designed for testing hypotheses rather than model selection, and thus suffer from the common drawbacks of hypothesis testing approaches. For instance, the type I error rate for the LRT increases with the number of comparisons (the multiple comparisons problem), and it is difficult to interpret the results of multiple models’ comparison that do not follow a nested structure. Choosing an approach for model selection is challenging in practice and remains an active area of ongoing research (e.g., Aho et al. (2014); Dziak et al. (2020)). In high-dimensional settings, bootstrap-based approaches (e.g., EIC or LRT) have been shown to perform best and are probably preferable when computationally feasible. For the case of the OU model, and when comparing models of different complexity (e.g., the OUM model with different optimum per trait), using a strategy combining multiple approaches may provide a more nuanced interpretation and a better understanding of the fitted models.

### The evolution of mammalian lower jaws and adaptation to diet regimes

Most methods of model selection support an Ornstein-Uhlenbeck process with two optima for the lower jaw evolution across metatherian and eutherian mammals. This result suggests that both groups of mammals (Metatheria and Eutheria) converged morphologically to fulfil the biomechanical demands of their specialised diets. The reconstructions of the mandible shape optimums for carnivorous and herbivorous species differ mainly in the corpus depth and the shape of the ramus, especially on the condylar and coronoid process (Fig. 5c). These differences are consistent with features associated with bite force, jaw movements, and stability in withstanding the stress resulting from food mastication (Hoshi 1971; Pérez-Barbería and Gordon 1999; Wroe and Milne 2007; Prevosti et al. 2012; Grossnickle 2020). The best supported model (OUM2), reveals a weak selection strength with a phylogenetic half-life about 1.1 (i.e., ∼ it takes 149 Ma to move halfway from the ancestral phenotype to the optimum for a 168 Ma phylogeny). However, it should be noted that, since the model assumes the selection strength is homogeneous across traits, this parameter describes the average strength of selection across the entire jaw, which may result in an underestimation of the selective forces acting on specific parts of the lower jaw. Allowing for trait-specific selection strengths would be possible but might be difficult to implement in practice due to the increasing number of parameters to optimize, which can become intractable in high-dimensional settings. As an alternative approach, it is possible to analyse the traits separately by anatomical modules or regions, when these can be identified *a priori* (e.g., developmental regions). This would allow to test whether the diet or some other selective factors act differently across structures (e.g., Goswami et al. 2019, 2022; Conith et al. 2022; Law et al. 2022). For instance, separating the jaw in two modules (an anterior module corresponding to the corpus, the tooth-bearing part of the mandible, and a posterior module corresponding to the ramus, where the masticatory muscles attach, Fig. S10, S11), we detected a stronger selection in the anterior than in the posterior region (see details in Supplementary material). The strong selective constraint in the anterior region is mainly related to the depth of the mandibular corpus, which is closely associated with mastication. In herbivorous species, the processing of tough materials (like plant fibres) and intensive chewing, tend to favour a deeper corpus than in carnivorous (Langenbach and Eijden 2001; Meloro et al. 2008). The posterior region (ramus) has clear biomechanical associations to food consumption, mainly as a surface for the anchoring of muscles involved in exerting bite force and mastication movements and stability through the jaw joint, or dissipation of torsional forces (Hoshi 1971; Biknevicius et al. 1996; Pérez-Barbería and Gordon 1999; Grossnickle 2020). However, the posterior region may be more functionally variable. This variation may result, for instance, from differences related to foraging strategy, prey size (in carnivores), or muscles performances (e.g., Spencer 1995; Meloro and O’Higgins 2011; Ito and Endo 2016), that can blur the form-function relationship (Fabre et al. 2018; Dickinson et al. 2021) or reBlect a many-to-one mapping traits, where distinct morphologies converge on similar performance or function (Wainwright 2005; Zelditch et al. 2017).

The OUM2 was selected by all the methods except the BIC, which selected an OU with a single optimum. It is widely acknowledged that the BIC penalises model selection more severely than the AIC, favouring simpler models with lower complexity (i.e., fewer estimated parameters). On the other hand, the AIC is known to sometimes favour more complex models, which may explain the incongruent selection between the two criteria. However, the consistent selection of the OUM2 model even when compared to models of larger or equal complexity (OUM4_cl_ and OUM2_dv_ respectively) suggests that overparameterisation may not be an issue here. Because the OUM2 model is supported by three out of four methods, including the more reliable EIC, we consider it to be the best option among the evaluated models. However, the discrepancy between the BIC and the other criteria, as well as the slight underfitting of the OUM2 to the data observed in the LRT, would suggest that our proposed OUM2 model might not fully capture some specificities of mammalian lower jaw evolution. For example, we evaluated the support for two hypotheses related to diet (i.e., OUM2 and OUM4_cl_) derived from an ancestral state reconstruction that places carnivory as the ancestral state in both placentals and marsupials, consistent with previous studies (Amador and Giannini 2021; Wu 2022). However, erroneous specifications of the diet categories, inaccuracies in the reconstructed states at the nodes of the phylogeny or in the location of regime shifts, or the omission of key lineages, would affect the inference and support of the OU models (Butler and King 2004; Ho and Ané 2014). The hypotheses evaluated here could be refined by increasing the number of species with characterised lower jaw morphologies to provide the necessary statistical power to analyse finer diet categorisation (i.e. different regimes for animal and plant-based diet), as for example those obtained from quantitative measures in (Grossnickle 2020), which would represent a more realistic picture of selective forces acting on the lower jaw morphology.

### Directions for future work

Several extensions of the Empirical Bayes framework can be envisioned. The first relates to the choice of the target matrix. Here, we proposed two commonly used target matrices (e.g., Schäfer and Strimmer 2005; Meyer 2011; Clavel et al. 2019; Ledoit and Wolf (2020) with various properties (e.g., trade-off between data structure complexity and requirements for rotation invariance; see discussion above). Although these two simple (diagonal) target matrices based on the sample covariance have shown good performance, they might be replaced by independent estimators that would effectively act as informative Bayesian priors. Conceptually, this simply requires selecting a different scale matrix in the Wishart distribution prior of equation (3). For instance, one could impose a given covariance structure between the traits based on some other empirical datasets (such as intraspecific covariances or G-matrix estimates for instance) or theoretical estimates (Clavel et al. 2019; Lam 2020; Ledoit and Wolf 2020). It may even be possible to use different target matrices, and therefore different priors, to compare specific hypotheses on the structure of the variance covariance matrix itself (Kolbe et al. 2011; Punzalan and Rowe 2016); the benefits, properties, and performance of this idea remain an open question for future research.

A second avenue for possible extensions is to relax the current constraint on the degrees of freedom of the *T* distribution. In the current implementation, we fixed *v* = *p* + 1, which then makes the prior on ***R*** more informative as *p* increases relative to *n*, and we demonstrated its efficiency. The performance of the regularised estimate ***R̃*** could, in principle, be improved by jointly optimising *v*. However, this is not an easy task. Some methods have been proposed but this parameter is generally difficult to estimate accurately (e.g., Ollila et al. 2020; Thompson et al. 2020; Pascal et al. 2021). Further investigations are required to assess the effectiveness and feasibility of these approaches in the context of phylogenetic comparative methods. The matrix-variate *T* distribution could also be used in Bayesian Markov chain Monte Carlo (MCMC) inference to Bit multivariate models with relaxed assumptions, such as branch-specific rates of evolution, at the same computational cost as simpler univariate models (given that the covariance matrix is explicitly integrated out).

A third potential area for improvements is model selection. For example, in the future it would be worth considering information criteria with higher-order corrections that have a lower computational cost than EIC. Such candidate criteria include the generalised information criterion (SGIC; (Konishi and Kitagawa 2003), or criteria that were developed for the small *n* large *p* problem in linear regressions (e.g., the Fisher information criterion FIC; Owrang and Jansson 2018) and its extended version (EFIC; Gohain and Jansson 2023).

Finally, regularised estimates have been shown to improve the performance of statistical approaches that rely on covariance matrices (e.g., Ullah and Jones 2015; Clavel and Morlon 2020). For instance, the PL-AR has demonstrated good performances in statistical analyses such as linear regressions and MANOVA (i.e., higher power and lower type I error rates) in comparison to other alternatives for high-dimensional data (Clavel and Morlon 2020). The performance of the Empirical Bayes approach in this type of analyses was not directly evaluated in this study, but we expect that it would perform similarly to those developed with the PL-AR regularisation in Clavel and Morlon (2020), while requiring less computational resources.

### Conclusions

The current massive generation of multivariate comparative datasets requires the development of efficient approaches to process and exploit these data. The proposed multivariate PCM uses the Empirical Bayes principle to address some of the current computational limitations when modelling high-dimensional phenotypic evolution on phylogenetic trees. The Empirical Bayes approach is accurate and fast, and because it can account for traits covariances without estimating or storing covariance matrices explicitly, it scales to very-high dimensional datasets (e.g. >4000 variables) more easily than previously proposed multivariate PCMs. In addition, it offers the possibility to obtain a regularised estimate of the evolutionary covariance matrix that can be used in many statistical approaches such as multivariate phylogenetic regressions and MANOVA tests. Model selection still remains a challenge to be addressed. The most effective approach identified here relies on the use of bootstrap resampling techniques, which are computationally intensive even though the Empirical bayes approach is fast. The properties and computational efficiency offered by the Empirical Bayes approach enable the extension of phylogenetic comparative methods to more complex and realistic models, and therefore the refinement of hypothesis testing, for high-dimensional comparative datasets.

## Acknowledgments

We are grateful to Nils Chabrol, Paul Bastide, Fabien Condamine and Julien Joseph for insightful discussions and constructive comments throughout the study. We also thank members of the J.C., H.M. and A.G. teams for their comments and feedback on the manuscript. Finally, we thank the two anonymous reviewers and the editors for their valuable feedback, which helped us to improve the manuscript. All the simulations were performed using the computing facilities of the CC-IN2P3/CNRS and LBBE/PRABI.

## Data availability statement

All the scripts necessary to reproduce the simulations and the empirical analyses on the mammalian data can be accessed through the Zenodo link https://doi.org/10.5281/zenodo.16380167. The approach (*mvgls()*, method=’EmpBayes’) and the functions used for model selection (*AIC()*, *BIC()*, *EIC()*, *LRT()*) are implemented in the R package ‘mvMORPH’ available on CRAN (https://cran.r-project.org/package=mvMORPH), and GitHub (https://github.com/JClavel/mvMORPH). Finally, the geometric morphometric data of the mammalian jaw, the phylogenetic tree and the ecological information used in the empirical analyses can be retrieved from the original publication supplementary material published on Dryad: https://doi.org/10.5061/dryad.b8gtht7c5.

