## Supplementary material for "An Empirical Bayes approach for the study of phenotypic evolution from high-dimensional data"

#### SUPPLEMENTARY METHODS AND DISCUSSION

##### 1. *Computing the confidence intervals*

We estimate the confidence intervals for the evolutionary parameters estimated by the empirical Bayes approach using the Fisher information matrix. This matrix describes the curvature of the likelihood surface around a given parameter $\hat{\theta}$ (Thacker 1989). Intuitively, a flat surface indicates that the uncertainty for the estimator is higher than for those with sharper likelihood surfaces (Ly et al. 2017). In practice, the Fisher Information matrix is often approximated using the second order derivative of the likelihood function evaluated at the MLE (the so-called Hessian or observed Fisher information matrix). The confidence intervals can then be estimated as (Efron and Hinkley 1978):

$$\hat{\theta} \pm cJ(\hat{\theta})^{-1/2}$$

Where $J(\hat{\theta})$ is the observed Fisher information matrix and $c$ is the corresponding critical value (e.g., $c$=1.96 to obtain a 95% confidence interval). We apply this formula to estimate the uncertainty around the evolutionary parameters describing the matrix $\boldsymbol{C}$ (i.e., $\alpha$ for the Ornstein-Uhlenbeck, $r$ for Early-Burst and $\lambda$ for Pagel's lambda models), by directly using the Hessian returned by the *optim()* function in the statistical software R. Note that this approach is not used for estimating uncertainty around the regularized estimate of the matrix $\boldsymbol{R}$.

We evaluated the performance of the proposed confidence intervals using simulations, following the general protocol described in the main text. Briefly, we simulated datasets for 100 lineages ($n = 100$) and six different $p$ traits ($p = c(20, 50, 80, 200, 500, 1000)$), for generating low and high-dimensional traits. We simulated a pure birth phylogenetic tree, scaled to unit height using the *pbtree()* function in the *phytools* package in R. We simulated $\boldsymbol{R}$ matrices using the two strategies described in the main text to generate *weakly-correlated* and *highly-correlated* traits (see *Methods: Testing the performances of the Empirical Bayes approach through simulations* in the main text for the complete description). We simulated data under three evolutionary models:

Early-Burst (r = -2.7); Ornstein-Uhlenbeck with one optimum ($\alpha$=2.7); and Pagel's lambda ($\lambda$=0.3). We fitted each model using the scaled-identity matrix as target. We obtained the Fisher information matrix for each one of the inferred parameters, directly from the *optim()* function in R, which is used for maximizing the likelihood in our Empirical Bayes approach. Then, we estimated the confidence intervals using the equation defined above. We repeated this procedure for every combination of parameters (*p*, the covariance matrix ***R***, and the model) and the coverage was quantified as the proportion of times that the true parameter fell within the confidence interval. To set a reference, we also estimated the same type of confidence intervals for models fit by the classical (multivariate normal distribution) maximum likelihood (ML) for the low dimensional simulated datasets. We fitted the models using ML with the *mvgls()* function from 'mvMORPH' package in R (Clavel et al. 2015).

The results, illustrated in the Figure S2, show the 95% coverage of the confidence intervals around the ML estimates (orange line). The coverage of the confidence intervals for our approach (blue line) depends on the structure of the traits. For *weakly-correlated* traits the coverage is always above 0.925 with slight changes as the number of variables *p* increases. For *highly-correlated* traits, as *p* increases the coverage decreases to almost 0.85 when *p* is 10 times *n*. Since confidence intervals are not regularly used in evolutionary biology and, to our knowledge, none of the few approaches modelling high-dimensional data include them, we have no references for comparison.

For high-dimensional datasets, the estimation of confidence intervals remains an area of active research. Other alternatives to estimate the confidence intervals, such as simulations (Boettiger et al. 2012), are possible. However, we prefer to use the Fisher information matrix rather than simulations-based approaches, since simulations of new datasets would require the use of the regularized covariance matrix ***R*** as input and it is not well understood how a regularized estimator can affect the output. Also, because obtaining the confidence intervals from the Fisher information matrix is simple and fast compared to simulations-based approaches.

The confidence intervals can be obtained from the *mvgls()* function in 'mvMORPH' package, through the argument '*FCI=TRUE*'.

#### 2. *Calculating the statistical loss of the regularized estimate of* ***R***

To compare the similarity between the simulated and the inferred ***R*** matrices, we used the Kullback-Leibler divergence given by (Kullback and Leibler 1951):

$$KL(\boldsymbol{R}||\widetilde{\boldsymbol{R}}) = \frac{1}{2}\left(tr\left(\widetilde{\boldsymbol{R}}^{-1}\boldsymbol{R}\right) - ln\frac{|\boldsymbol{R}|}{|\widetilde{\boldsymbol{R}}|} + p\right) \quad \text{(S1)}$$

And the quadratic loss given by (Van Wieringen and Peeters 2016; Clavel et al. 2019):

$$QL(\widetilde{\boldsymbol{R}}, \boldsymbol{R}) = \left\|\widetilde{\boldsymbol{R}}^{-1}\boldsymbol{R} - \boldsymbol{I}_p\right\|^2 \quad \text{(S2)}$$

Where $\boldsymbol{R}$ is the true covariance matrix, $\widetilde{\boldsymbol{R}}$ the regularized estimate obtained by the various approaches, and $p$ stands for the number of traits.

3. *Efficient estimation of the determinant in the likelihood function of the Matrix-variate T distribution*

Estimating the likelihood function for the matrix -variate $T$ distribution (equation 4 in main text) require the calculation of the determinant of a matrix whose dimension is either $n$ x $n$ or $p$ x $p$ (Gupta and Nagar 2000) (Theorem 4.3.3):

$$|\boldsymbol{I}_n + \boldsymbol{C}^{-1}(\boldsymbol{Y} - \boldsymbol{\Theta})\boldsymbol{\Psi}^{-1}(\boldsymbol{Y} - \boldsymbol{\Theta})^T|^{-\frac{1}{2}(v+n+p-1)} = \left|\boldsymbol{I}_p + \boldsymbol{\Psi}^{-1}(\boldsymbol{Y} - \boldsymbol{\Theta})^T\boldsymbol{C}^{-1}(\boldsymbol{Y} - \boldsymbol{\Theta})\right|^{-\frac{1}{2}(v+n+p-1)}$$

This equivalence can be exploited to estimate the determinant on the matrix with the lowest dimension. Similarly, because $\boldsymbol{\Psi}$ is a diagonal matrix, multiplication by its inverse in the formula above, as well as the determinant $|\boldsymbol{\Psi}|^{-\frac{1}{2}n}$ in equation (4) can be computed very efficiently. Finally, estimation of the $\boldsymbol{C}^{-1}$ matrix and its determinant can be computed efficiently using a pruning-like algorithm as implemented in the *pruning()* function in 'mvMORPH' (see (Stone 2011; Khabbazian et al. 2016)).

4. *Evaluating morphological convergences in the jaw shape in mammals by anatomical modules*

To evaluate whether selection acts differently across the mandibular regions, we fitted the evolutionary models to two anatomical modules. We divided the lower jaw into i) *anterior region*, composed mainly by the mandibular corpus, and ii) *posterior region* or ramus and condylar process (Figure S10). Rather than aligning each dataset separately, we split the aligned coordinates (for the whole lower jaw) into the two modules to facilitate the comparison with the analyses on the complete lower jaw in the main text. Then, we followed the same procedure as for the complete lower jaw (see *Methods* in the main text), to evaluate differences in morphology between 1) carnivorous and herbivorous lineages (OUM2); 2) carnivorous and herbivorous lineages within each clade, Eutheria and Metatheria ($OUM4_{cl}$); and 3) Eutheria and Metatheria, irrespective of their diet regime ($OUM2_{dv}$). Following the analyses on the whole lower jaw, we also fit the Ornstein-Uhlenbeck (OU) model with a single optimum, the Early-Burst (EB), and Brownian motion (BM) model on both modules. We tested the support for each models using the AIC, BIC and EIC (1000 bootstraps). Furthermore, we compared the support between the OU models with LRTs (using 1000 bootstrap samples for generating each distribution).

We found the same results for both regions than for the complete jaws, with the AIC and EIC supporting the OUM2 as the best model, while the BIC supported the OU with a single optimum (Table S3). Likewise, the LRTs favoured the OUM2 over the other models in both modules (Table S4). The OUM2 models describe a different selection strength between the two modules. The selection strength in the anterior module is higher than for the posterior module and the complete jaw, with an $\alpha \sim 0.011$, that corresponds to 2.6 half-lives (or ~63 Mya relative to the ~168 Mya of the phylogenetic tree). In contrast, in the posterior module we inferred a lower selection strength than for the anterior module and the complete jaw, with $\alpha \sim 0.0038$, corresponding to a low half-life (<1 or ~ 182 Mya). The reconstructions of the optimal morphologies are congruent with the morphologies inferred using the complete dataset (Figure S11).

Analysing the anatomical modules separately helped in understanding how the mammalian jaw has evolved. The results suggest that the two anatomical modules evolved towards the herbivorous and carnivorous optimum with different strength (see *Discussion* section in the main text). The estimated selection strength for the entire jaw is intermediate to the strength estimated for the anterior and posterior modules (see Table 2), thus emphasising the conclusion that the $\alpha$ parameter summarises the strength of selection across the whole structure.

**SUPPLEMENTARY TABLES**

| *Parameter* | *Description* |
|---|---|
| $n$ | Number of lineages |
| $p$ | Number of traits |
| $\boldsymbol{V}$ | Variance-covariance matrix for the vectorized dataset **Y**. The matrix is of dimension $np \times np$. Under some assumptions, it is possible to decompose it as $\boldsymbol{V} = \boldsymbol{C} \otimes \boldsymbol{R}$, and then describe it as a matrix-variate distribution (e.g., the matrix variate normal or the matrix variate $T$ distributions) |
| $\boldsymbol{\Theta}$ | Mean of a matrix-variate distribution. Matrix of dimensions $n \times p$. Its vectorised version $vec(\boldsymbol{\Theta})$, of dimension $np$, describes the mean for a multivariate normal distribution. |
| $\boldsymbol{C}$ | Variance-covariance matrix of dimensions $n \times n$. It corresponds to the phylogenetic variance- covariance matrix in this study |
| $\boldsymbol{R}$ | Variance-covariance matrix of dimensions $p \times p$ of a matrix-variate distribution. It corresponds to the variance- covariance matrix for the traits |
| $\widehat{\boldsymbol{R}}$ | Maximum likelihood estimate of $\boldsymbol{R}$ |
| $\widetilde{\boldsymbol{R}}$ | Empirical Bayes (regularised) estimate of $\boldsymbol{R}$ |
| $\boldsymbol{\Psi}$ | Scale matrix parameter of the matrix-variate $T$ distribution. It corresponds to the target matrix in the regularization. |
| $\kappa$ | Degrees of freedom of the Inverse Wishart distribution. **Fixed** to $v + p - 1$ |
| $v$ | Degrees of freedom of the matrix-variate $T$ distribution. **Fixed** to $p + 1$ |
| $\boldsymbol{\beta}$ | Ancestral states of each trait. Vector of dimension $p$ or $p \times m$, when using the OUM model. |
| $\widehat{\boldsymbol{\beta}}$ | Maximum likelihood estimates for the ancestral states of each trait. **Estimated from equation (2).** |
| $\boldsymbol{X}$ | Design matrix. Matrix of ones. The number of columns corresponds to the number of fixed effects (1 for all the models except OUM, where the number of columns $m$ corresponds to the number of selective regimes). Note that in regression settings, this matrix corresponds to the fixed predictors. |

Table S1. Summary of all the parameters used throughout the main text.

| Approach | *p* | Total time (s) | | Total memory (GB) | | Max Mem by expression (bytes) | | Number of Expressions |
|---|---|---|---|---|---|---|---|---|
| | | mean | SD | mean | SD | mean | SD | |
| EmpBayes | 100 | 0.099 | 0.047 | 0.038 | 0.011 | 8847.5 | 2533.585 | 4.00E+04 |
| EmpBayes(R) | | 0.102 | 0.025 | 0.040 | 0.011 | 9401.88 | 2543.681 | 8.01E+04 |
| PL-AR | | 0.270 | 0.116 | 0.069 | 0.022 | 25834.44 | 8308.737 | 8.01E+04 |
| PL-AR(LOOCV) | | 2.170 | 0.740 | 1.718 | 0.569 | 101360.04 | 33440.46 | 8.01E+04 |
| PL-QR(LOOCV) | | 7.005 | 1.150 | 3.502 | 0.247 | 142960.88 | 10037.294 | 8.01E+04 |
| EmpBayes | 200 | 0.116 | 0.039 | 0.053 | 0.007 | 8713.12 | 1098.165 | 8.00E+04 |
| EmpBayes(R) | | 0.145 | 0.042 | 0.06 | 0.007 | 9275.12 | 1098.165 | 3.20E+05 |
| PL-AR | | 0.673 | 0.183 | 0.19 | 0.017 | 23571.4 | 2031.813 | 3.20E+05 |
| PL-AR(LOOCV) | | 4.746 | 1.101 | 5.425 | 0.487 | 90725.8 | 8067.309 | 3.20E+05 |
| PL-QR(LOOCV) | | 25.983 | 9.080 | 14.496 | 3.122 | 158888 | 34044.76 | 3.20E+05 |
| EmpBayes | 1000 | 0.135 | 0.048 | 0.136 | 0.002 | 9496.8 | 85.118 | 4.00E+05 |
| EmpBayes(R) | | 0.728 | 0.245 | 0.257 | 0.002 | 10058.8 | 85.118 | 8.00E+06 |
| PL-AR | | 5.203 | 1.758 | 3.546 | 0.125 | 24745 | 756.349 | 8.00E+06 |
| PL-AR(LOOCV) | | 84.902 | 23.510 | 111.32 | 4.071 | 85873 | 2993.326 | 8.00E+06 |
| PL-QR(LOOCV) | | 1551.94 | 639.663 | 475.366 | 96.327 | 223471.84 | 44448.511 | 8.00E+06 |
| EmpBayes | 2000 | 0.396 | 0.121 | 0.443 | 0.08 | 17354.84 | 1710.494 | 8.00E+05 |
| EmpBayes(R) | | 2.911 | 0.527 | 0.922 | 0.08 | 17916.84 | 1710.494 | 3.20E+07 |
| PL-AR | | 35.076 | 9.89 | 18.426 | 4.368 | 35820 | 6752.903 | 3.20E+07 |
| EmpBayes | 4000 | 1.133 | 0.171 | 1.415 | 0.141 | 31517.84 | 1534.549 | 1.60E+06 |
| EmpBayes(R) | | 19.167 | 2.131 | 3.326 | 0.141 | 32079.84 | 1534.549 | 1.28E+08 |
| PL-AR | | 207.349 | 65.29 | 82.396 | 23.626 | 47553.36 | 9228.211 | 1.28E+08 |

Table S2. Computational performance summary statistics for the Empirical Bayes and Penalized-likelihood (PL) approaches compared in the main text. The table summarises the results of 50 simulations with trees of $n$=50 species across a range of traits dimensions ($p$). The total time and total memory allocated in the statistical software R, are reported in seconds and gigabytes (GB), respectively. The maximum memory used by the most expensive expression (Max Mem by expression) is shown in bytes. Also, the total number of expressions used when running each approach is shown. Approaches: EmpBayes: Empirical Bayes; EmpBayes(R): EmpBayes computing the matrix $\widetilde{\boldsymbol{R}}$ explicitly; PL-AR: PL with the archetypal ridge penalty and using the fast leaving-one-out-cross-validation (LOOCV) algorithm (H&L). PL-AR(LOOCV): PL-AR using the regular leaving-one-out-cross-validation; PL-QR(LOOCV): PL with a quadratic ridge penalty using a regular LOOCV.

| | Model | ΔAIC | ΔBIC | ΔEIC | SE EIC | α |
|---|---|---|---|---|---|---|
| ***Anterior module*** | *BM* | -447.669 | -259.997 | -448.877 | 9.004 | - |
| | *EB* | -449.669 | -264.552 | -446.838 | 8.610 | - |
| | *OU* | -185.117 | **0.000** | -218.677 | 8.843 | 0.0105 (0.0081, 0.0129) |
| | *OUM2* | **0.000** | -151.993 | **0.000** | 8.849 | 0.011 (0.0085, 0.0135) |
| | $OUM2^{dv}$ | -711.419 | -1537.637 | -691.048 | 8.366 | 0 |
| | $OUM2_{cl}$ | -581.107 | -733.101 | -586.108 | 8.750 | 0.0032 (0, 0.0030) |
| ***Posterior module*** | *BM* | -325.417 | -128.128 | -363.050 | 17.449 | - |
| | *EB* | -327.417 | -132.682 | -432.203 | 18.187 | - |
| | *OU* | -194.734 | **0.000** | -229.0989 | 18.310 | 0.0036 (0.0022, 0.0050) |
| | *OUM2* | **0.000** | -341.579 | **0.000** | 17.223 | 0.0039 (0.0024, 0.0053) |
| | $OUM2^{dv}$ | -689.584 | -2103.793 | -657.619 | 16.656 | 0 |
| | $OUM2_{cl}$ | -646.317 | -987.897 | -785.413 | 18.275 | 0 |

Table S3. Support for the models fit on the anterior and posterior modules of the mammal jaws, using different information criteria. The difference relative to the lowest value (delta) is reported for AIC, BIC and EIC. The standard error (SE) of the estimated EIC is also reported. The alpha parameter (selection strength) for the OU models is shown, with the confidence intervals within parentheses. Because the lower bound estimated for the $OUM2_{cl}$ is negative (-0.0013), the value is set to zero. See Table 2 in main text. BM: Brownian motion; OU: Ornstein-Uhlenbeck with one optimum; EB: Early-Burst; OUM: Ornstein-Uhlenbeck with multiple optima following the ecologies OUM2: herbivorous/carnivorous, $OUM2_{dv}$: Ornstein-Uhlenbeck with multiple optima following the clades: Metatheria/Eutheria, and $OUM4_{cl}$: herbivorous Methateria/Eutheria and carnivorous Methateria/Eutheria. In bold the model supported as the most likely is highlighted.

| | Null model | Alternative model | $LR_{obs}$ | p-value |
|---|---|---|---|---|
| **Anterior module** | OU | OUM2 | 449.1179 | **0.001** |
| | OU | $OUM2_{dv}$ | -131.9897 | 0.888 |
| | OUM2 | $OUM2_{dv}$ | -593.1774 | 0.448 |
| | OUM2 | $OUM4_{cl}$ | -183.4197 | 0.994 |
| **Posterior module** | OU | OUM2 | 614.7343 | **0** |
| | OU | $OUM2_{dv}$ | -31.58342 | 0.879 |
| | OUM2 | $OUM2_{dv}$ | -643.7331 | 0.331 |
| | OUM2 | $OUM4_{cl}$ | 150.4151 | 0.884 |

Table S4. Results of the likelihood ratio tests (LRT) for the comparisons between the various Ornstein-Uhlenbeck models (OU), for the anterior and posterior modules of the mammal jaws. The null hypothesis is rejected when the p-value is under the critical value of 0.05 (highlighted in bold). $LR_{obs}$ stands for the observed likelihood ratio.

**SUPPLEMENTARY FIGURES**

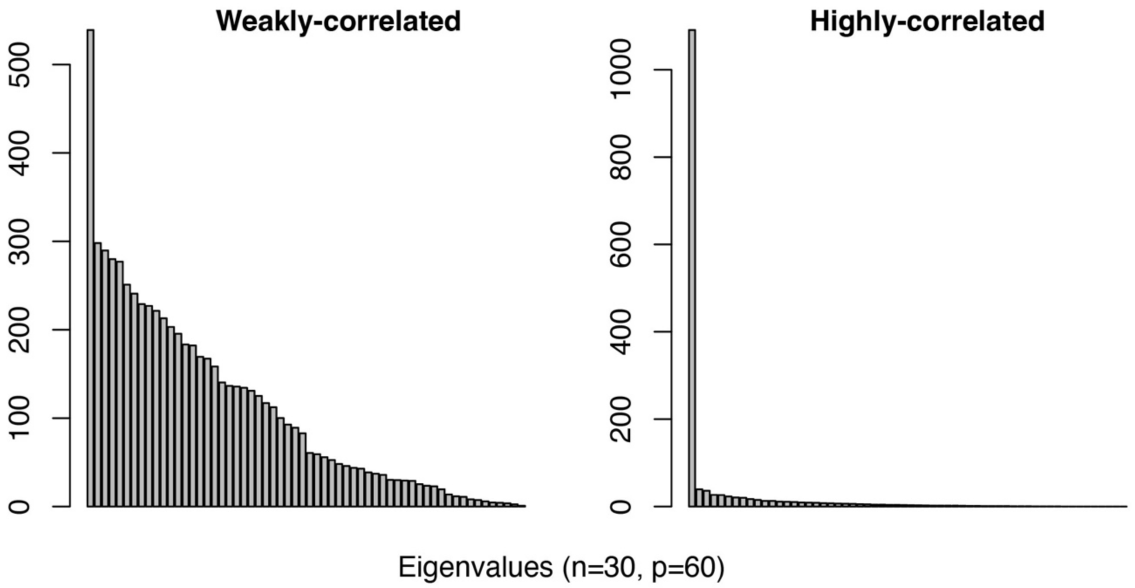


Figure S1. Eigenvalues distribution for the covariance matrices simulated with either the *weakly-correlated* and *highly-correlated* approaches (see the *Methods* in the main text for a detailed description).

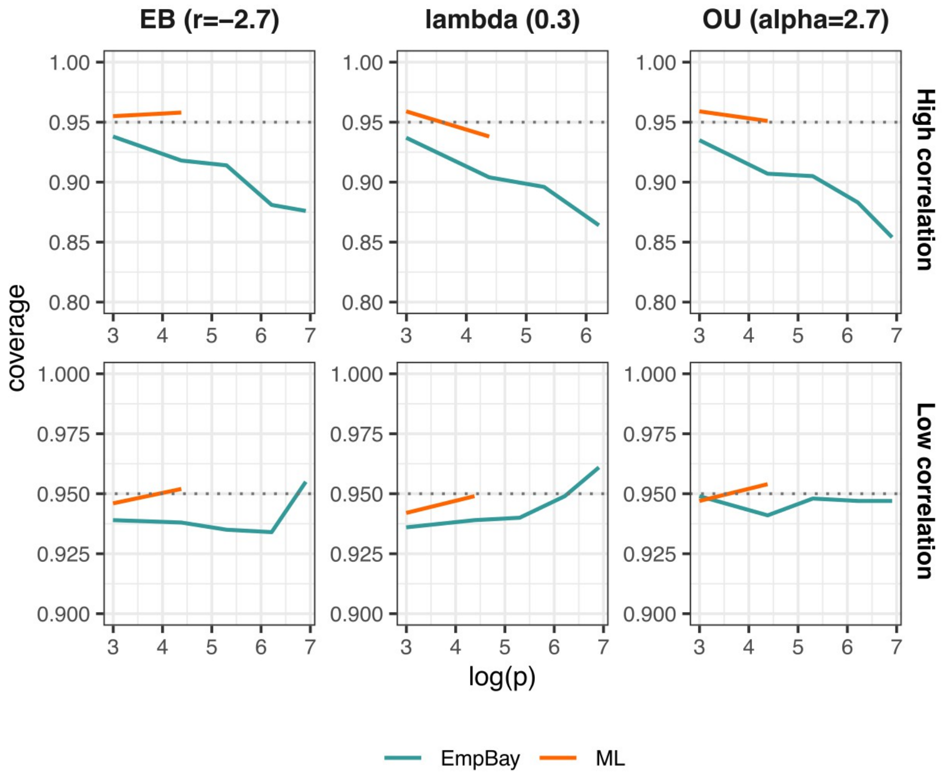


Figure S2. Coverage probability of the confidence intervals constructed using the Fisher information matrix for parameters inferred by ML (orange line) and the Empirical bayes (blue line) approach, when $n$ = 100 and $p$ varies from 20 to 1000. The dotted line indicates the expected coverage for 95% confidence intervals. The confidence intervals of the parameters inferred by the Empirical Bayes approach are around 94% coverage when the traits are *weakly-correlated* (lower panel), while the coverage decreases as $p$ increases with values down to around 85% when *p=1000* in *highly-correlated* datasets. The confidence intervals for the MLE of conventional comparative methods were estimated for the low dimensional datasets only because these approaches are not applicable when $p<n$. Models: EB: Early Burst; lambda: Pagel's lambda; OU: Ornstein-Uhlenbeck with one optimum. The parameter value used for the simulations is indicated between parentheses.

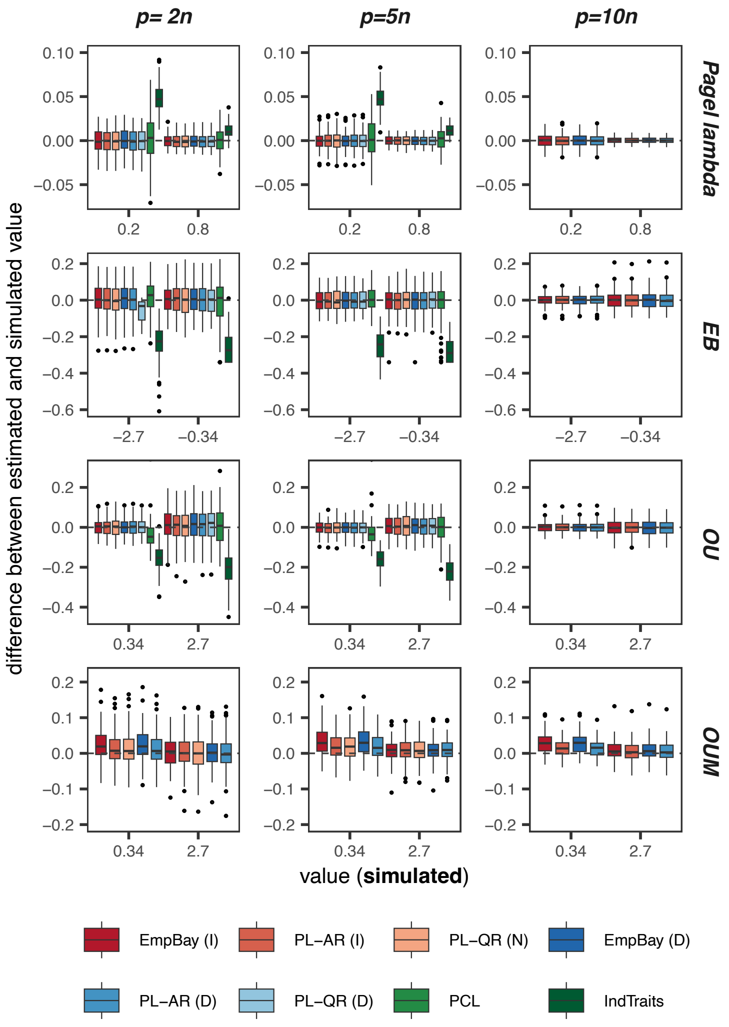


Figure S3. Difference between the true parameter used for the simulations (traits *weakly-correlated*) and the parameter estimated and for four evolutionary models: Pagel's lambda, Early Burst (EB), Ornstein-Uhlenbeck with one (OU) and multiple optima (OUM). The dashed line indicates no differences (value = 0). Boxplots represent the distribution based on 100 simulations. From the left to the right, the ratio between the number of lineages (*n=100*) and the number of variables (*p*) increases. Approaches: EmpBay: Empirical Bayes; PL-AR: Penalized likelihood – Archetypal Ridge; PL-QR:Penalized likelihood – Quadratic Ridge; PCL: Pairwise composite likelihood; IndTraits: joint estimation over the individual likelihoods for each traits (this correspond to multivariate models where the covariance matrix is forced to be diagonal – i.e., the traits are assumed to be independent). (I), (N) and (D), stands for identity, null, and diagonal target matrix, respectively. The PL-QR approaches were not evaluated for *p=10n* because of their computationally prohibitive cost.

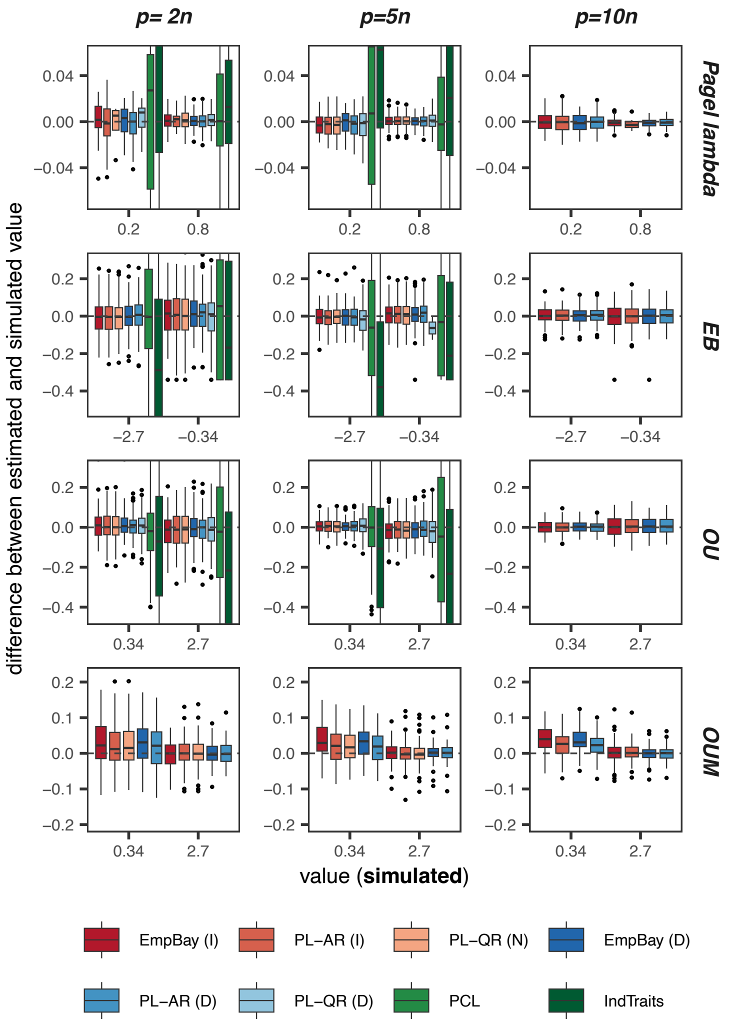


Figure S4. Difference between the true parameter used for the simulations (traits *highly-correlated*) and the parameter estimated for four evolutionary models: Pagel's lambda, Early Burst (EB), Ornstein-Uhlenbeck with one (OU) and multiple optima (OUM). The traits were simulated using covariance matrices generated with a pronounced skewness of the eigenvalues (see the main text for details). Boxplot represent the distribution based on 100 simulations. From the left to the right, the ratio between the number of lineages (*n=100*) and the number of variables (*p*) increases. For the approaches based on regularization, three target matrices were considered: identity (I), null (N; only available for the PL-QR approach), and diagonal matrix (D). Approaches: EmpBay: Empirical Bayes; PL-AR: Penalized likelihood – Archetypal Ridge; PL-QR: Penalized likelihood – Quadratic Ridge; PCL: Pairwise composite likelihood; IndTraits: joint estimation over the individual likelihoods for each traits (this correspond to multivariate models where the covariance matrix is forced to be diagonal – i.e., the traits are assumed to be independent). The PL-QR approaches were not evaluated for *p=10n* because of their computationally prohibitive cost.

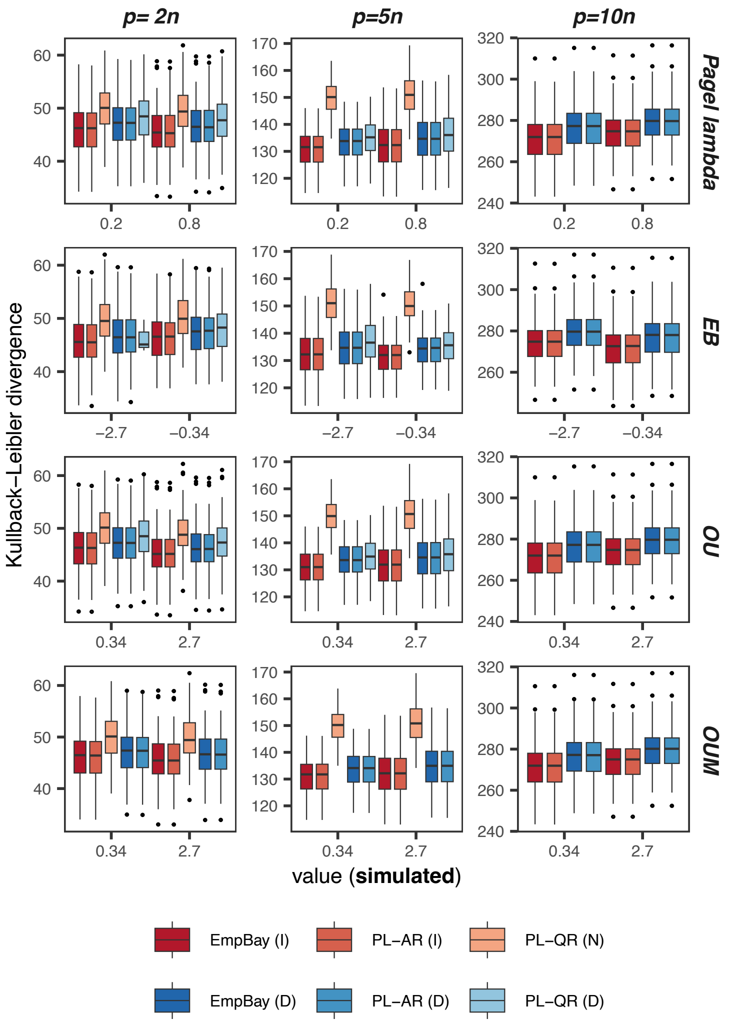


Figure S5. Kullback-Leibler divergence between the estimated and simulated traits covariance matrix under Pagel's lambda model, Early Burst (EB), Ornstein-Uhlenbeck with one (OU) and multiple optima (OUM), for traits *weakly-correlated*. Boxplot represent the distribution based on 100 simulations. From the left to the right, the ratio between the number of lineages (*n=100*) and the number of variables (*p*) increases. For the approaches based on regularization, three target matrices were considered: identity (I), null (N; only available for the PL-QR approach), and diagonal matrix (D). Approaches: EmpBay: Empirical Bayes; PL-AR: Penalized likelihood – Archetypal Ridge; PL-QR: Penalized likelihood – Quadratic Ridge; PCL: Pairwise composite likelihood; IndTraits: joint estimation over the individual likelihoods for each traits (this correspond to multivariate models where the covariance matrix is forced to be diagonal – i.e., the traits are assumed to be independent). The PL-QR approaches were not evaluated for *p=10n* because of their computationally prohibitive cost.

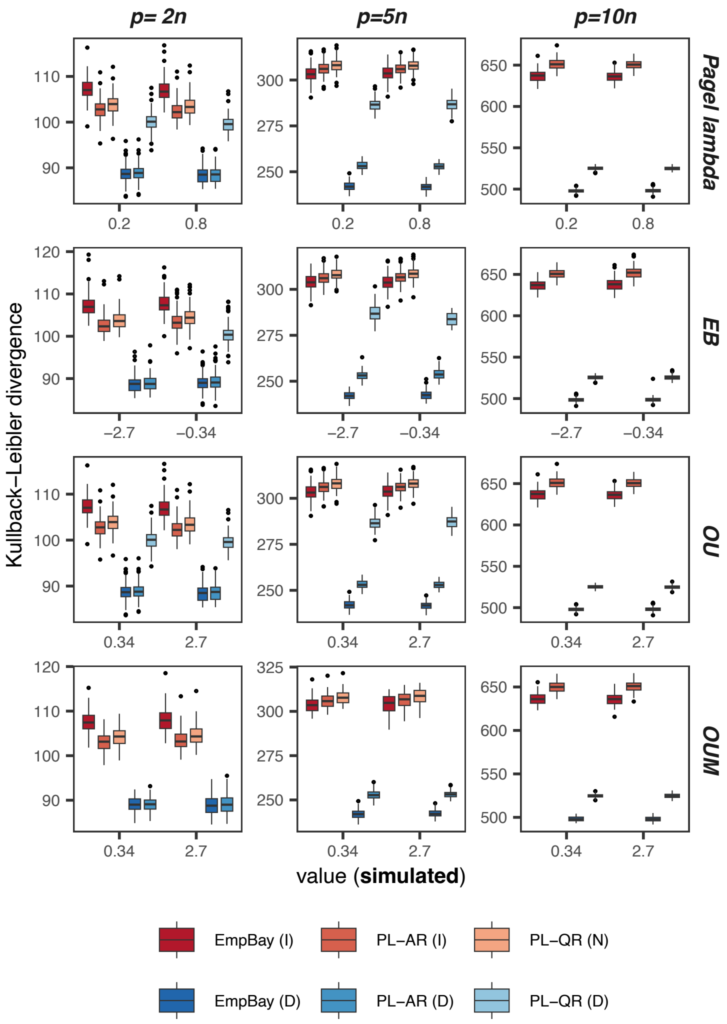


Figure S6. Kullback-Leibler divergence between the estimated and simulated traits covariance matrix under Pagel's lambda model, Early Burst (EB), Ornstein-Uhlenbeck with one (OU) and multiple optima (OUM), for traits *highly-correlated* (i.e., pronounced skewness of the eigenvalues). Boxplot represent the distribution based on 100 simulations. From the left to the right, the ratio between the number of lineages (*n=100*) and the number of variables (*p*) increases. For the approaches based on regularization, three target matrices were considered: identity (I), null (N; only available for the PL-QR approach), and diagonal matrix (D). Approaches: EmpBay: Empirical Bayes; PL-AR: Penalized likelihood – Archetypal Ridge; PL-QR: Penalized likelihood – Quadratic Ridge; PCL: Pairwise composite likelihood; IndTraits: joint estimation over the individual likelihoods for each traits (this correspond to multivariate models where the covariance matrix is forced to be diagonal – i.e., the traits are assumed to be independent). The PL-QR approaches were not evaluated for *p=10n* because of their computationally prohibitive cost.

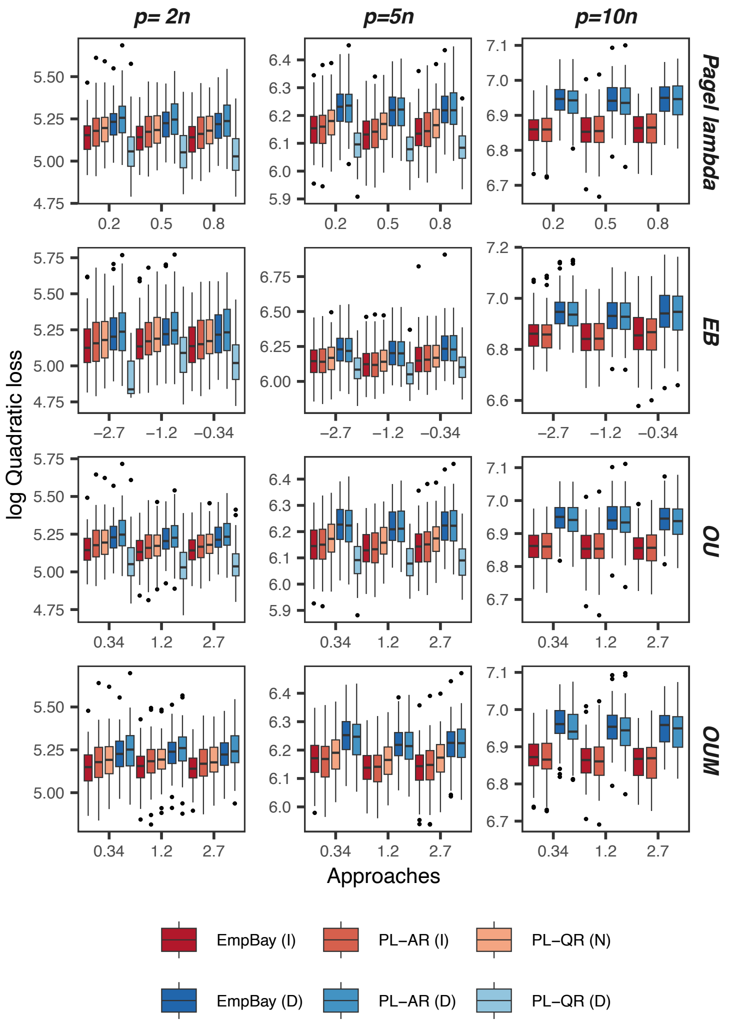


Figure S7. Quadratic loss (log) between the simulated and estimated traits covariance matrix under Pagel's lambda, Early Burst (EB), Ornstein-Uhlenbeck with one (OU) and multiple optima (OUM), when the traits *weakly-correlated*. Boxplot represent the distribution based on 100 simulations. From the left to the right, the ratio between the number of lineages (*n=100*) and the number of variables (*p*) increases. For the approaches based on regularization, three target matrices were considered: identity (I), null (N; only available for the PL-QR approach), and diagonal matrix (D). Approaches: EmpBay: Empirical Bayes; PL-AR: Penalized likelihood – Archetypal Ridge; PL-QR: Penalized likelihood – Quadratic Ridge; PCL: Pairwise composite likelihood; IndTraits: joint estimation over the individual likelihoods for each traits (this correspond to multivariate models where the covariance matrix is forced to be diagonal – i.e., the traits are assumed to be independent). The PL-QR approaches were not evaluated for *p=10n* because of their computationally prohibitive cost.

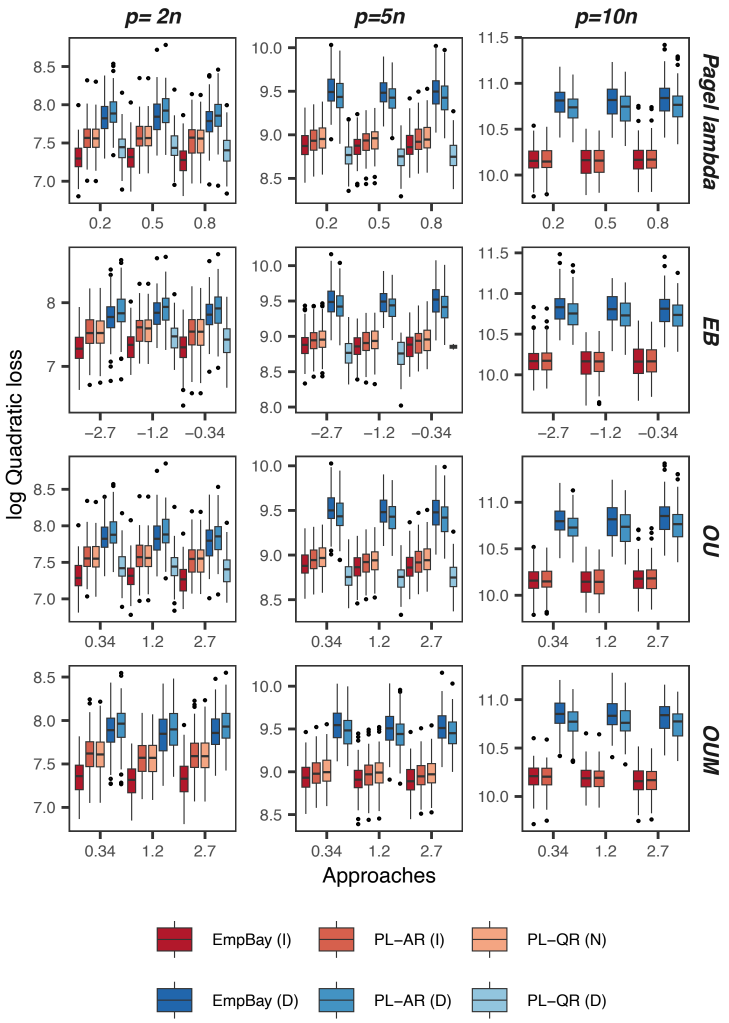


Figure S8. Quadratic loss (log) between the simulated and estimated traits covariance matrix under Pagel's lambda, Early Burst, Ornstein-Uhlenbeck with one (OU) and multiple optima (OUM), when the *highly-correlated* traits (i.e., pronounced skewness of the eigenvalues). Boxplot represent the distribution based on 100 simulations. From the left to the right, the ratio between the number of lineages (*n=100*) and the number of variables (*p*) increases. For the approaches based on regularization, three target matrices were considered: identity (I), null (N; only available for the PL-QR approach) and diagonal matrix (D). Approaches: EmpBay: Empirical Bayes; PL-AR: Penalized likelihood – Archetypal Ridge; PL-QR: Penalized likelihood – Quadratic Ridge; PCL: Pairwise composite likelihood; IndTraits: joint estimation over the individual likelihoods for each traits (this correspond to multivariate models where the covariance matrix is forced to be diagonal – i.e., the traits are assumed to be independent). The PL-QR approaches were not evaluated for *p=10n* because of their computationally prohibitive cost.

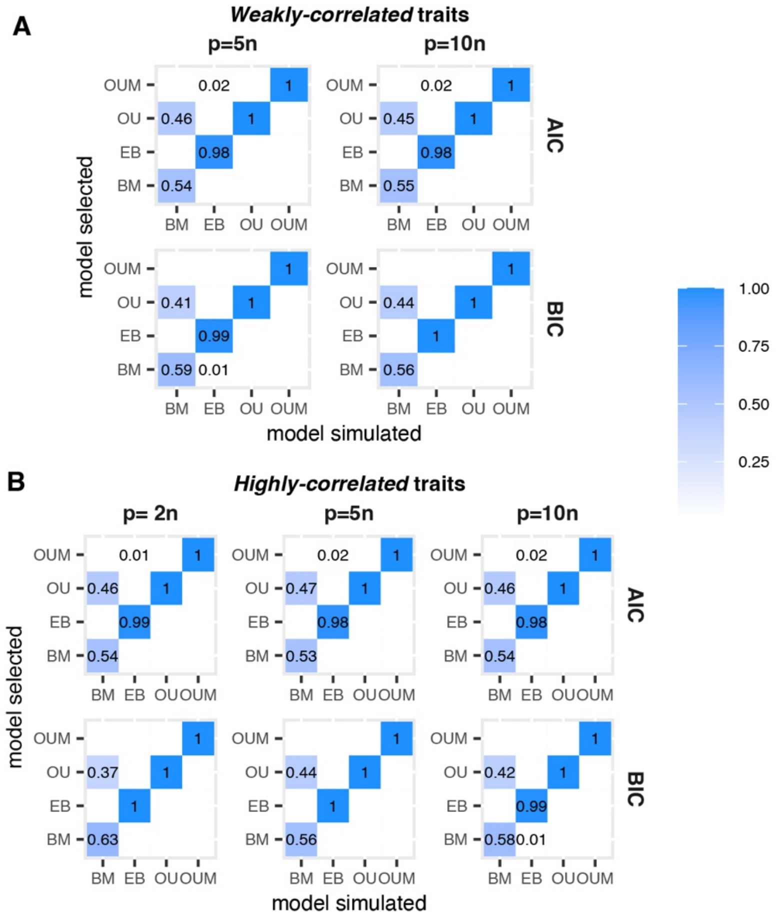


Figure S9. Proportion of times the simulated model is selected (lowest information criterion value) using the Akaike (AIC), and the Bayesian (BIC) Information criteria, across 100 simulations. In panel **A**) the traits were simulated as *weakly-correlated* and in **B**) they were simulated as *highly-correlated*. In panel **A**, *p=2n* is omited as it is already illustrated in the main text (Figure 3). In all the simulations, the number of species was *n*=100. Both, AIC and BIC show high consistency for all the models evaluated except BM. Models: Brownian Motion (BM), Early Burst (EB), Ornstein-Uhlenbeck with one (OU) and multiple optima (OUM).

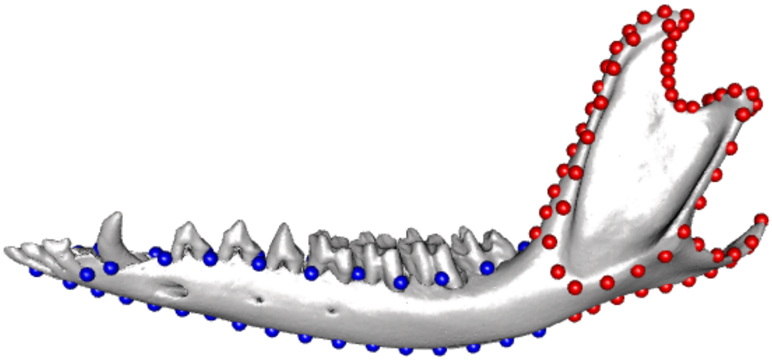

Figure S10. Positions of the landmarks and semi-landmarks for the anterior (blue) and posterior (red) modules of the lower mammal jaw. The raw coordinates and mesh of the northern brown bandicoot (*Isoodon macrourus*) were used for the illustration.

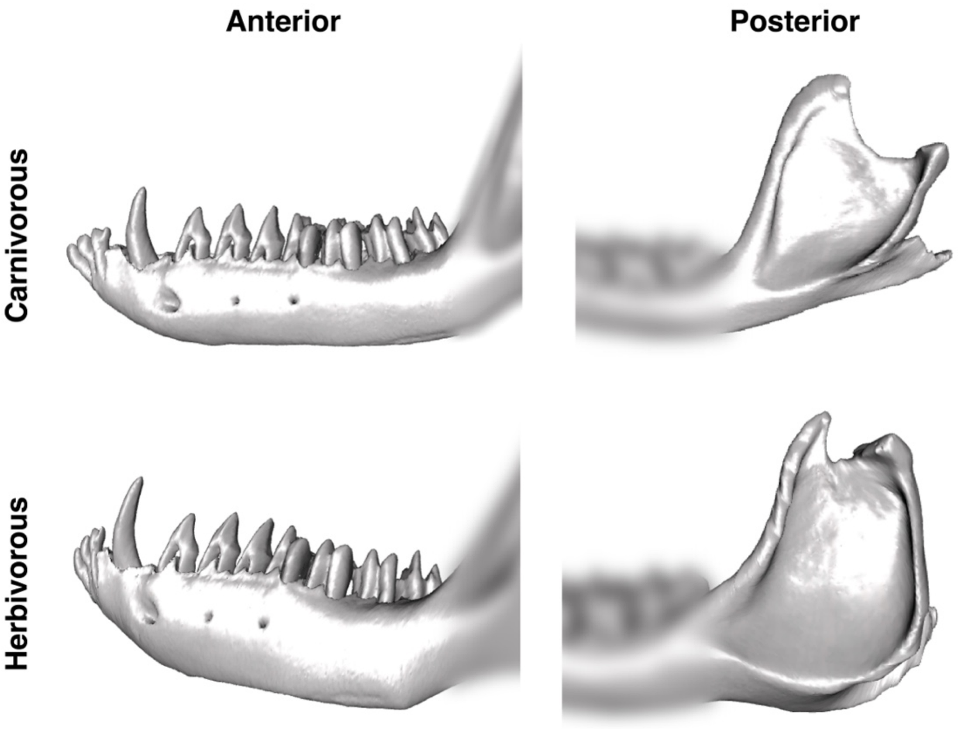


Figure S11. Reconstruction of the optima for each dietary regime based on the analyses of the two anatomical modules. Reconstructions were obtained by warping the mesh of the northern brown bandicoot (*Isoodon macrourus*) onto the optimum coordinates using thin-plate spline interpolation.

### Example_Empirical_Bayes_mvMORPH

Paola Montoya

2026-03-03

#### Running the Empirical Bayes approach in mvgls()

```r
set.seed(2807)

library(mvMORPH)
```

##### Loading mvMORPH and data

```
## Loading required package: phytools

## Loading required package: ape

## Loading required package: maps

## Loading required package: corpcor

## Loading required package: subplex

## ##
## ## mvMORPH package (1.2.2) beta version 04/04/25

##  # On est déjà grands :  . . . . . . . . . . ##
##  ##                ,--.  ,--.                ##
##  ##               /   | /    \               ##
##  ##               `|  ||  ()  |              ##
##  ##                |  | \    /               ##
##  ##                `--'  `--'                ##
##  ##               _ | ___ | _                ##
##  ##             |             |              ##
##  ##         + - - - - - - - - - - +          ##
##  ##         |                     |          ##
##  ##     + - - - - - - - - - - - - - - +      ##
##  ##                                          ##
##  ##             __  __  ___  ___ ___ _  _    ##
##  ##    _ ____ _|  \/  |/ _ \| _ \ _ \ || |   ##
##  ##   | '  \ V / |\/| | (_) |   /  _/ __ |   ##
##  ##   |_|_|_\_/|_|  |_|\___/|_|_\_| |_||_|   ##
##  ## . . .  ... mais, ce n’est que le début :) #
##

##

## ## Multivariate evolutionary models

## ##
## ## See the tutorials: browseVignettes("mvMORPH")
```

```
## ##
## ## To cite package 'mvMORPH': citation("mvMORPH")
## ##
```

```
data("phyllostomid")
dim(phyllostomid$mandible)
```

```
## [1] 49 73
```

phyllostomid is a dataset of 72 2D geometric morphometric coordinates for 49 species, describing the lower jaw of phyllostomid bats. It was published by Montero et al. 2011

```
# Phylogenetic tree
phy = phyllostomid$tree

# Traits
traits = phyllostomid$mandible

# Selecting only traits corresponding to geometric morphometric coordinates
traits = traits[,-1]
```

##### Fitting the evolutionary models using the Empirical Bayes approach

The Empirical Bayes approach allows fitting different trait evolution models, by setting method='EmpBayes' in the mvgls() function. Unlike the other approaches in the function, it does not require a penalty method. Currently, two targets are available: 'unitVariance' (the default option) and 'Variance'. The FCI argument specifies the computation of the confidence intervals using the Fisher information matrix. By default, FCI=FALSE. See ?mvgls() for an explanation of the other arguments.

```
# Ornstein-Uhlenbeck with a single optimum (OU)
fit.ou = mvgls(traits~1,tree=phy,model='OU',method='EmpBayes',target='unitVariance',
               error=TRUE,FCI=TRUE)

# Brownian motion (BM)
fit.bm = mvgls(traits~1,tree=phy,model='BM',method='EmpBayes',target='unitVariance',
               error=TRUE,FCI=TRUE)

# Early-Burst (EB)
fit.eb = mvgls(traits~1,tree=phy,model='EB',method='EmpBayes',target='unitVariance',
               error=TRUE,FCI=TRUE)
```

##### Exploring the output

Once the model is fitted, different parameters describing the model are obtained.

```

# evolutionary parameter r for EB model
fit.eb$param
```

**Parameters:**

```
##      Var2
## -2.231392
```

**Confidence intervals** The confidence intervals are computed for the parameters describing the trait evolution model, other than the regularised estimate of the covariance matrix R (e.g., alpha in OU, r in EB, or lambda in Pagel's lambda). lw and up stand for lower and upper interval respectively.

```r
fit.eb$FCI
```

```
##   lw.Var2   up.Var2
## -3.250640 -1.212144
```

```r
fit.eb$mserr
```

**Error estimated**

```
##      Var3
## 0.1695043
```

**Regularised estimate for R** Note that the regularised estimate for R is p x p dimensions

```r
dim(fit.eb$sigma$Pinv)
```

```
## [1] 72 72
```

```r
head(fit.eb$sigma$Pinv)
```

```
##              x1           y1           x2           y2           x3
## x1 2.717301e-04 1.225303e-05 2.775020e-04 3.243706e-05 1.058528e-04
## y1 1.225303e-05 2.884248e-04 8.243184e-05 1.673094e-04 6.125082e-05
## x2 2.775020e-04 8.243184e-05 5.170753e-04 5.131623e-05 2.040072e-04
## y2 3.243706e-05 1.673094e-04 5.131623e-05 1.780653e-04 3.585802e-05
## x3 1.058528e-04 6.125082e-05 2.040072e-04 3.585802e-05 2.987824e-04
## y3 5.623388e-05 6.213715e-05 6.600104e-05 6.965853e-05 9.563630e-06
##              y3            x4            y4            x5            y5
## x1 5.623388e-05 -6.014295e-05 -5.863912e-06 -5.445898e-05 -8.862248e-06
## y1 6.213715e-05  1.072751e-04  1.561315e-05  6.632729e-05  5.877817e-06
## x2 6.600104e-05 -1.328130e-04  3.687143e-05 -1.345212e-04  2.982826e-05
## y2 6.965853e-05  9.278682e-05 -7.334864e-06  6.890004e-05 -1.837056e-05
## x3 9.563630e-06 -1.623820e-04  3.487573e-05 -1.565821e-04  6.562597e-06
## y3 9.970649e-05  2.866412e-05  1.868389e-05  2.149815e-05  9.351602e-06
##               x6            y6            x7           y7            x8
## x1 -4.223069e-05  9.513334e-06 -4.315365e-05 4.969237e-05 -5.097164e-05
## y1  2.474754e-05  1.993011e-05 -6.306820e-06 3.618615e-05 -3.152501e-05
## x2 -1.162086e-04  5.594807e-05 -1.019975e-04 1.047922e-04 -9.468529e-05
## y2  4.062245e-05 -4.030358e-06  1.648895e-05 1.687342e-05 -7.781857e-06
## x3 -1.295293e-04  8.345752e-07 -1.042328e-04 6.023419e-06 -8.888617e-05
## y3  1.341245e-05  1.843293e-05  5.331632e-06 3.395933e-05 -6.488369e-06
##              y8            x9           y9           x10          y10
## x1 9.530677e-05 -4.937870e-05 1.375853e-04 -3.157478e-05 1.717414e-04
## y1 4.322608e-05 -6.676155e-05 4.238565e-05 -9.788176e-05 2.517641e-05
## x2 1.539224e-04 -8.064158e-05 1.960081e-04 -4.943133e-05 2.246873e-04
## y2 3.415961e-05 -3.946372e-05 4.678181e-05 -6.489045e-05 4.660479e-05
## x3 1.455189e-05 -7.854211e-05 2.366155e-05 -6.375810e-05 2.823358e-05
## y3 5.058520e-05 -1.914313e-05 6.388138e-05 -2.557386e-05 6.891386e-05
##              x11          y11           x12          y12           x13
## x1 -1.557259e-06 1.821339e-04  2.228308e-06 1.558228e-04 -1.029153e-05
## y1 -1.199235e-04 7.240174e-06 -1.005505e-04 1.171421e-05 -8.976445e-05
## x2 -1.154400e-06 2.249213e-04  1.566272e-05 2.039320e-04  1.977627e-06
## y2 -7.971894e-05 3.948219e-05 -6.583902e-05 3.939017e-05 -6.444545e-05
## x3 -3.908146e-05 2.417873e-05 -1.802679e-05 2.569680e-05 -2.558034e-05
## y3 -2.614280e-05 6.781706e-05 -1.490481e-05 6.230946e-05 -1.776970e-05
```

```
##              y13           x14          y14           x15          y15
## x1 1.302510e-04 -3.677535e-06 9.054664e-05 -4.765003e-07 4.419405e-05
## y1 2.524734e-05 -6.092127e-05 4.034580e-05 -2.442568e-05 4.300702e-05
## x2 1.838472e-04  1.044067e-05 1.439988e-04  2.672318e-05 8.624420e-05
## y2 3.911788e-05 -5.482904e-05 3.576979e-05 -3.822299e-05 2.838526e-05
## x3 2.126654e-05 -1.130097e-05 1.193711e-05  1.762057e-05 2.224706e-06
## y3 5.482296e-05 -1.596712e-05 4.074674e-05 -1.373946e-05 2.052693e-05
##               x16          y16           x17           y17           x18
## x1  4.553203e-06 2.155777e-05  2.020095e-05  2.650969e-05  1.969260e-05
## y1  4.321032e-06 3.353151e-05  1.754161e-05  2.159603e-05  1.513042e-05
## x2  5.363864e-05 4.322302e-05  9.384069e-05  2.268114e-05  8.993412e-05
## y2 -2.320550e-05 1.739002e-05 -9.011684e-06  1.389014e-05 -1.161428e-05
## x3  4.387372e-05 3.596986e-06  5.620561e-05  4.862341e-06  7.330361e-05
## y3 -1.383553e-05 3.364441e-07 -2.399057e-06 -6.040258e-06 -6.808344e-06
##               y18           x19           y19           x20           y20
## x1  9.605755e-06  2.847600e-05 -2.088175e-06  3.935034e-05 -1.303149e-05
## y1  2.695916e-05  2.430614e-05  2.418800e-05  2.615965e-05  2.016859e-05
## x2 -6.066880e-07  9.095880e-05 -1.249224e-05  9.325165e-05 -2.641574e-05
## y2  1.562530e-05 -4.648865e-06  1.257638e-05 -2.174747e-06  1.147229e-05
## x3  1.661660e-06  9.372404e-05  1.065678e-06  1.095921e-04  5.856656e-06
## y3 -6.605377e-06 -2.489728e-06 -7.844660e-06  1.568670e-06 -9.648710e-06
##               x21           y21           x22           y22           x23
## x1  4.814651e-05 -2.971618e-05  2.638561e-05 -5.684536e-05  2.573581e-06
## y1  2.270470e-05  1.628701e-05  3.412127e-05  2.747049e-05  3.499110e-05
## x2  9.014847e-05 -4.895423e-05  4.088396e-05 -8.554047e-05 -5.016407e-06
## y2 -6.110015e-07  9.974944e-06  7.769022e-06  7.772689e-06  1.501492e-05
## x3  1.141140e-04  1.462821e-05  8.818460e-05  1.256813e-05  6.194052e-05
## y3  3.430906e-06 -1.356460e-05 -1.600219e-05 -1.626241e-05 -1.607135e-05
##               y23           x24           y24           x25           y25
## x1 -8.629237e-05 -9.735851e-06 -1.339307e-04 -2.385619e-05 -1.475567e-04
## y1 -1.109801e-05  1.323091e-05 -3.154529e-05 -1.656307e-05 -2.865978e-05
## x2 -1.155404e-04 -3.669288e-05 -1.631079e-04 -7.371585e-05 -1.840867e-04
## y2 -1.938405e-05  7.796316e-06 -2.627523e-05 -4.278518e-06 -2.054504e-05
## x3  2.992658e-05  4.014935e-05  6.425039e-06  1.363485e-05 -4.395322e-05
## y3 -3.667186e-05 -1.356475e-05 -4.825348e-05 -1.462851e-05 -4.431446e-05
##               x26           y26           x27           y27           x28
## x1 -5.181674e-05 -1.372143e-04 -7.949482e-05 -1.102817e-04 -9.338692e-05
## y1 -5.059643e-05 -4.129934e-05 -6.144229e-05 -9.255272e-05 -3.210736e-05
## x2 -1.344963e-04 -1.910350e-04 -1.861333e-04 -1.617527e-04 -2.057384e-04
## y2 -1.711750e-05 -3.095597e-05 -1.626431e-05 -7.356983e-05  4.425494e-06
## x3 -3.023924e-05 -5.778970e-05 -7.391015e-05 -4.974400e-05 -1.113010e-04
## y3 -2.072323e-05 -4.698445e-05 -2.278354e-05 -5.618233e-05 -1.443553e-05
##               y28           x29           y29           x30           y30
## x1 -9.837197e-05 -9.914966e-05 -8.526069e-05 -9.429714e-05 -6.920017e-05
## y1 -1.292859e-04  3.434727e-06 -1.506972e-04  2.841720e-05 -1.490006e-04
## x2 -1.409211e-04 -2.048877e-04 -1.231953e-04 -1.816274e-04 -1.057599e-04
## y2 -1.086943e-04  2.503068e-05 -1.249963e-04  3.604081e-05 -1.198860e-04
## x3 -4.178794e-05 -1.292619e-04 -3.293675e-05 -1.132754e-04 -2.647778e-05
## y3 -6.338473e-05 -5.359376e-06 -6.464255e-05  7.971313e-08 -6.035507e-05
##               x31           y31           x32           y32           x33
## x1 -7.829212e-05 -5.605979e-05 -5.287620e-05 -4.979227e-05 -1.935294e-05
## y1  3.814640e-05 -1.319057e-04  3.640714e-05 -1.071881e-04  3.240655e-05
## x2 -1.367000e-04 -9.609425e-05 -7.297360e-05 -9.530176e-05  4.786221e-06
## y2  3.513767e-05 -1.013704e-04  2.529149e-05 -7.804292e-05  1.397088e-05
```

```
## x3 -6.596856e-05 -3.191048e-05 -4.127617e-06 -4.085276e-05  4.594842e-05
## y3  1.664446e-06 -5.105154e-05  1.402842e-06 -4.396714e-05  4.349703e-06
##              y33          x34           y34            x35           y35
## x1 -4.479026e-05 1.725620e-05 -4.106569e-05  4.885648e-05 -2.778303e-05
## y1 -7.655412e-05 3.046840e-05 -4.659653e-05  2.925253e-05 -1.171889e-05
## x2 -1.003497e-04 8.580812e-05 -1.045111e-04  1.440114e-04 -8.935683e-05
## y2 -5.232655e-05 5.368731e-06 -2.840657e-05 -1.374189e-08 -6.546202e-06
## x3 -4.499063e-05 5.998152e-05 -3.655820e-05  3.134565e-05 -6.742330e-06
## y3 -3.957147e-05 1.354766e-05 -3.815046e-05  2.707951e-05 -3.604079e-05
##              x36           y36
## x1  5.561402e-05 -2.137766e-05
## y1  1.344424e-05  2.232584e-05
## x2  1.270300e-04 -6.563255e-05
## y2 -1.012397e-05  8.681009e-06
## x3 -3.402898e-05  2.643287e-05
## y3  3.500204e-05 -3.408780e-05
```

```
fit.eb$logLik
```

###### Log-likelihood

```
## [1] 13141.93
```

###### Using mulitple information criteria for model selection

Given that the Empirical Bayes approach only works with the Restricted Maximum Likelihood (REML), for model comparison we propose to estimate the Maximum Likelihood (ML) using the REML estimates. For this, it is needed to set REML=FALSE.

```
aic.ou = AIC(fit.ou,REML=FALSE)
aic.bm = AIC(fit.bm,REML=FALSE)
aic.eb = AIC(fit.eb,REML=FALSE)
which.min(c('OU'=aic.ou$AIC,'BM'=aic.bm$AIC,'EB'=aic.eb$AIC))
```

###### Akaike information criteria

```
## BM.Var1
##       2
```

```
bic.ou = BIC(fit.ou,REML=FALSE)
bic.bm = BIC(fit.bm,REML=FALSE)
bic.eb = BIC(fit.eb,REML=FALSE)
which.min(c('OU'=bic.ou$BIC,'BM'=bic.bm$BIC,'EB'=bic.eb$BIC))
```

###### Bayesian information criteria

```
## BM.Var1
##       2
```

**Fitting an Ornstein-Uhlenbeck with multiple optima (OUM)** Using the Empirical Bayes approach, it is also possible to fit the OUM model to estimate the strength of selection by selective regimes (categories). These selective regimes are represented by a discrete variable mapped onto the phylogenetic tree. The map then includes the state of the regime (the discrete variable) at the tips and nodes (reconstructions). In this example, we will use different categories describing the dietary resources consumed by phyllostomid bats.

```
diet2 = as.character(phyllostomid$grp1)
# Note that every element should be named according to the tip labels

# Evaluating the best model for the reconstruction of the ancestral states
c('sym'=AIC(ace(diet2, phy, type = 'discrete', model="SYM", marginal = TRUE)),
  'er'=AIC(ace(diet2, phy, type = 'discrete', model="ER", marginal = TRUE)),
  'ard'=AIC(ace(diet2, phy, type = 'discrete', model="ARD", marginal = TRUE)))
```

**H1: 2 optima according to the diet: High-mastication Low-mastication**

```
##      sym       er      ard
## 29.56446 29.56446 29.69548
```

```
# Reconstruction of ancestral states
anc_rec.d2 = ace(diet2, phy, type = "discrete", model="SYM", marginal = TRUE)
diet_tree2 = mapping.asr(phy, anc_rec.d2, diet2)
colors = c("#4575b4","#35978f")
names(colors)=c('High-mastication','Low-mastication')

# Visualizing the final hypothesis
plot(diet_tree2,col=colors,fsize=0.5,ftype='i')
```

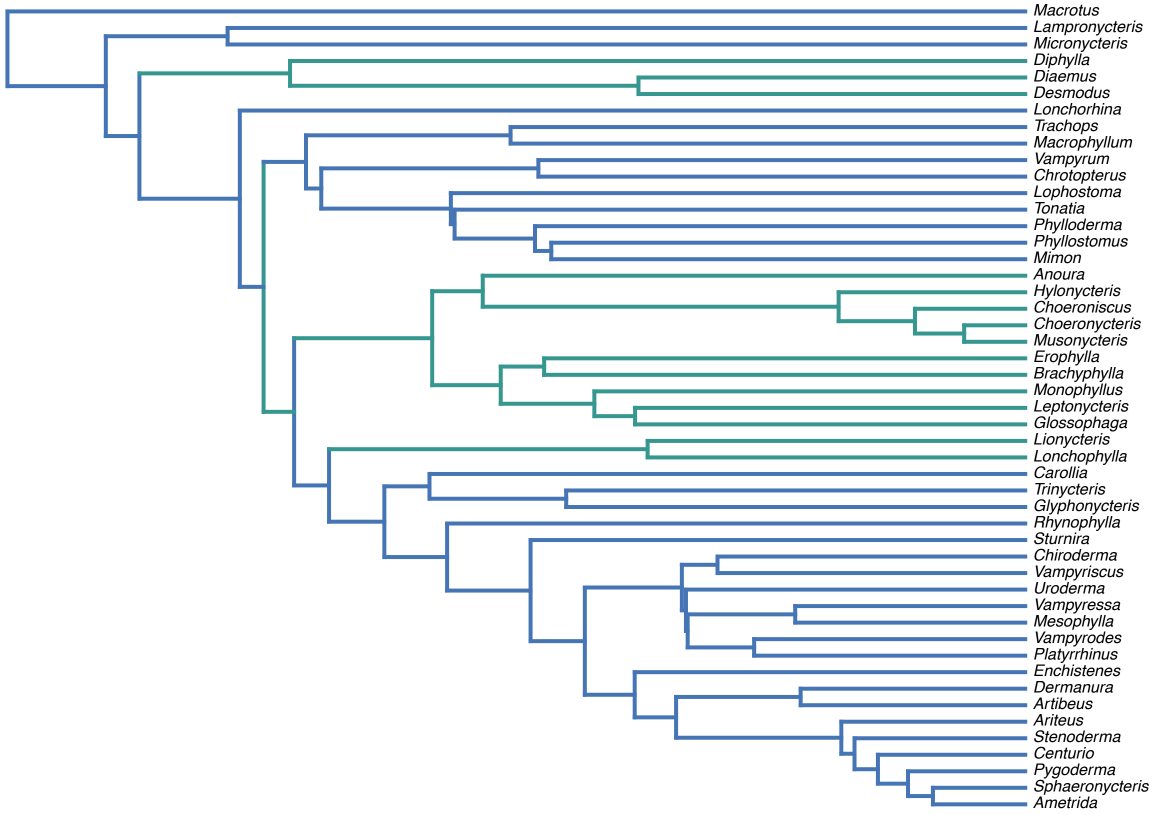


```
diet4 = as.character(phyllostomid$grp3)
# Note that every element should be named according to the tip labels
```

```r
# Evaluating the best model for the reconstruction of the ancestral states
c('sym'=AIC(ace(diet4, phy, type = 'discrete', model="SYM", marginal = TRUE)),
  'er'=AIC(ace(diet4, phy, type = 'discrete', model="ER", marginal = TRUE)),
  'ard'=AIC(ace(diet4, phy, type = 'discrete', model="ARD", marginal = TRUE)))
```

**H2: 4 optima according to the diet: Frugivory, Animalivory, Nectarivory, Sanguivory**

```
##      sym       er      ard
## 59.10674 51.78294 67.95145
```

```r
# Reconstruction of ancestral states
anc_rec.d4 = ace(diet4, phy, type = "discrete", model="ER", marginal = TRUE)
diet_tree4 = mapping.asr(phy, anc_rec.d4, diet4)
colors = c('#7b3294','#c2a5cf','#a6dba0','#008837')
names(colors)=c('Frugivory','Animalivory', 'Nectarivory', 'Sanguivory')

# Visualizing the final hypothesis
plot(diet_tree4,col=colors,fsize=0.5,ftype='i')
```

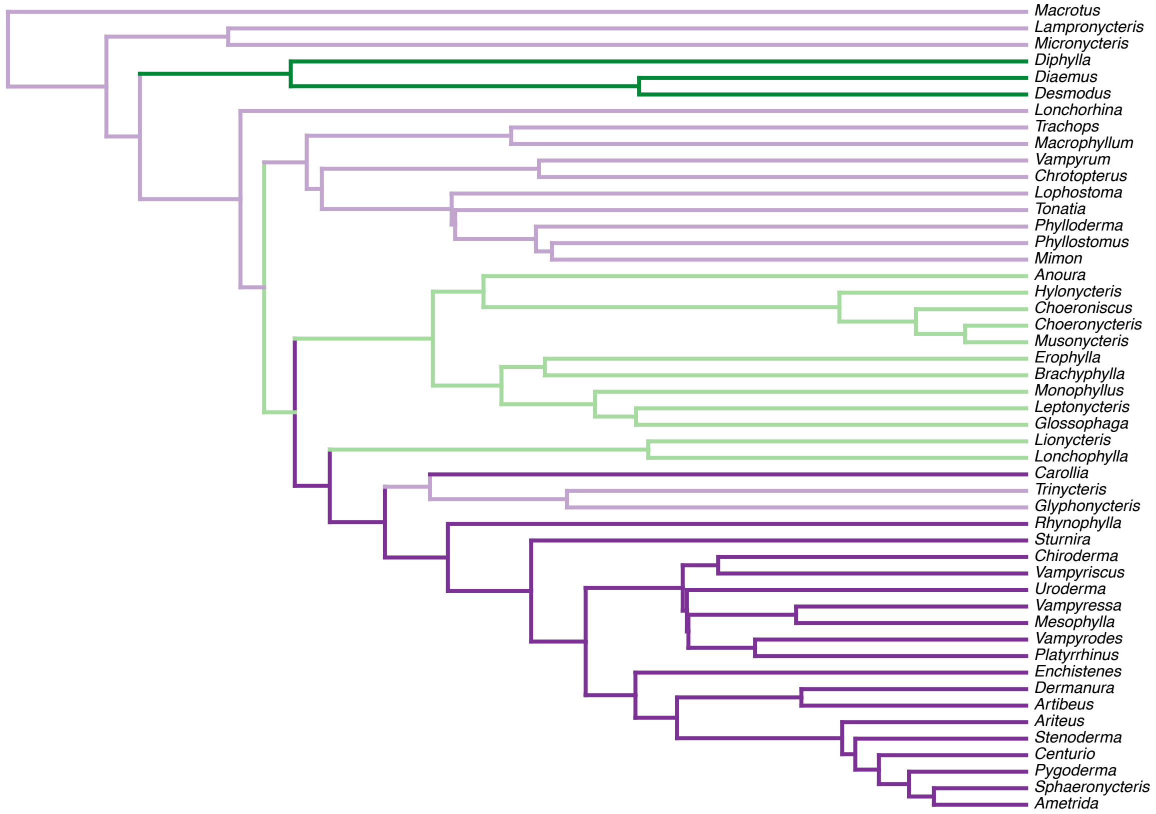


```r
# Ornstein-Uhlenbeck with multiple optima (OUM)
fit.oum.2 = mvgls(traits~diet2,tree=diet_tree2,model='OUM',method='EmpBayes',
                  target='unitVariance',error=TRUE,FCI=TRUE)

fit.oum.4 = mvgls(traits~diet4,tree=diet_tree4,model='OUM',method='EmpBayes',
                  target='unitVariance',error=TRUE,FCI=TRUE)

# Parameters: alpha values
```

```r
c('OUM2'=fit.oum.2$param,'OUM4'=fit.oum.4$param)
```

**Fitting the model**

```
##    OUM2.Var2    OUM4.Var2
## 0.0000000001 1.5069512000
```

```r
# Confidence intervals for H2
c('OUM4'=fit.oum.4$param,fit.oum.4$FCI[1],
  fit.oum.4$FCI[2])
```

```
## OUM4.Var2   lw.Var2   up.Var2
## 1.5069512 0.2368269 2.7770755
```

```r
##### Extended information criteria
eic.bm = EIC(fit.bm,REML=FALSE,nboot=100)
eic.oum4 = EIC(fit.oum.4,REML=FALSE,nboot=100)
which.min(c('BM'=eic.bm$EIC,'OUM.H2'=eic.oum4$EIC))
```

**Model selection**

```
## OUM.H2.Var1
##           2
```
